# Saracatinib, a Selective Src Kinase Inhibitor, Blocks Fibrotic Responses in *In Vitro, In Vivo* and *Ex Vivo* Models of Pulmonary Fibrosis

**DOI:** 10.1101/2022.01.04.474955

**Authors:** Farida Ahangari, Christine Becker, Daniel G. Foster, Maurizio Chioccioli, Meghan Nelson, Keriann Beke, Xing Wang, Benjamin Readhead, Carly Meador, Kelly Correll, Loukia Lili, Helen M. Roybal, Kadi-Ann Rose, Shuizi Ding, Thomas Barnthaler, Natalie Briones, Giuseppe Deluliis, Jonas C. Schupp, Qin Li, Norihito Omote, Yael Aschner, Katrina W. Kopf, Björn Magnusson, Ryan Hicks, Anna Backmark, Leslie P. Cousens, Joel T. Dudley, Naftali Kaminski, Gregory P. Downey

## Abstract

Idiopathic Pulmonary Fibrosis (IPF) is a chronic, progressive, and often fatal disorder. Two FDA approved anti-fibrotic drugs, nintedanib and pirfenidone, slow the rate of decline in lung function, but responses are variable and side effects are common. Using an *in-silico* data-driven approach, we identified a robust connection between the transcriptomic perturbations in IPF disease and those induced by saracatinib, a selective Src kinase inhibitor, originally developed for oncological indications. Based on these observations, we hypothesized that saracatinib would be effective at attenuating pulmonary fibrosis. We investigated the anti-fibrotic efficacy of saracatinib relative to nintedanib and pirfenidone in three preclinical models: (i) *in vitro* in normal human lung fibroblasts (NHLFs); (ii) *in vivo* in bleomycin and recombinant adenovirus transforming growth factor-beta (Ad-TGF-β) murine models of pulmonary fibrosis; and (iii) *ex vivo* in precision cut lung slices from these mouse models. In each model, the effectiveness of saracatinib in blocking fibrogenic responses was equal or superior to nintedanib and pirfenidone.

## Introduction

Idiopathic pulmonary fibrosis (IPF) is a chronic, relentless, and ultimately fatal disorder characterized by progressive scarring (fibrosis) of the lung parenchyma (1-3). The median survival for IPF patients is three years from diagnosis with most patients dying from respiratory failure due to disease progression (4). The overall prevalence of IPF worldwide and in United States ranges from 3.3-45.1, and 14.0 - 63.0 cases per 100,000 respectively*(4-6)*. Recent studies suggest that both the prevalence of, and mortality from, IPF are increasing *(4,7)*.

The mechanisms driving pulmonary fibrosis remain incompletely understood and both genetic and environmental factors appear to be important in disease pathogenesis *(8,9)*. A widely accepted hypothesis is that, in genetically susceptible individuals, the lung is repeatedly injured (via an unknown cause) and aberrant repair provokes release and activation of fibrogenic mediators, myofibroblast accumulation, and deposition of excess extracellular matrix (ECM) resulting in progressive fibrosis *(10-12)*. Cellular responses to profibrotic mediators are frequently transduced through transmembrane receptors via intracellular signaling pathways that are controlled in part by Src family tyrosine kinases (SFK) that include Src, Yes, Fyn, Fgr, Lck, Hck, Blk, Lyn, and Frk *(13,14)*. SFK are involved in a range of signaling pathways essential for cellular homeostasis such as proliferation, differentiation, motility, adhesion, and cytoskeletal organization. Studies in preclinical models of IPF suggest that several Src-dependent processes contribute to IPF pathogenesis, including myofibroblast differentiation and fibrogenic gene expression *(15,16)*.

Treatment of IPF remains suboptimal. Two anti-fibrotic drugs, pirfenidone (Esbriet^®^) and nintedanib (OFEV^®^), were approved by the FDA in 2014 for the treatment of IPF *(17-19)*. Clinical trials and real-world experience demonstrate that, while on average, both drugs slow the rate of decline in lung function, responses are variable, and these drugs neither cure IPF nor improve the major symptoms including cough and dyspnea *(20,21)*. Thus, there remains an urgent need for development of more effective therapies that safely modify the course of IPF and restore quality of life.

An evolving understanding of the molecular underpinnings of human diseases has provided opportunities to precisely target disease-specific pathways using bioinformatic methods to mine genomic, molecular, and clinical data *(22-24)*. This approach has been used successfully to identify connections between disease and drug ‘molecular signatures’, revealing opportunities to use existing drugs in new therapeutic areas (‘computational drug repurposing’) *(22-25)*. As part of a data-driven approach to repurpose Phase-II ready compounds for new diseases *(26)*, we identified saracatinib as a potential therapeutic compound for IPF. Saracatinib is a potent and selective Src kinase inhibitor, originally developed for oncological indications *(27-29)*. Building on the initial insights gleaned from transcriptomic connections, and the evidence strongly supporting a pivotal role for Src kinase in IPF pathophysiology, we sought to determine the anti-fibrotic efficacy of saracatinib relative to nintedanib and pirfenidone in pre-clinical cell culture and animal models of IPF and to elucidate the molecular signatures of pathological fibrogenesis and drug responsiveness.

## Results

### Computational drug re-purposing approach identifies a connection between saracatinib and IPF

At the outset, we undertook a disease agnostic, data-driven approach to explore novel connections between diseases and compounds previously tested in clinical studies. A set of 32 compounds was selected for this analysis based on a combination of factors that included experience in Phase 1 and Phase 2 clinical studies and potential for further clinical development beyond the disease for which the compound was originally designed (‘drug repositioning’) (https://openinnovation.astrazeneca.com/data-library.html#transcriptomicprofilingdata) *(26)*. The method applied was based on a modified connectivity mapping approach *(26, 31, 32)* where the transcriptomic signature of each compound was compared computationally with transcriptomic signatures of human diseases. Briefly, differential gene expression signatures for each compound were generated by preforming RNA sequencing on two different cell lines (A549 and MCF7) after exposure to two concentrations of each of the 32 compounds (e.g., “Compound-A A549 High dose”, “Compound-A MCF7 High dose”, “Compound-A A549 Low dose”, “Compound-A MCF7 Low dose”). This analysis was done blinded to the compound identity or chemistry and was generated using the genome-wide pattern of mRNA changes in cell-matched compound-versus vehicle-treated samples. (**Figure 1A**). A disease transcriptomic library, consisting of over 700 unique disease signatures, was built from publicly available gene expression data *(23,33)*. To identify novel clinical indications for each compound, a “connectivity score” was calculated by comparing each disease signature to each compound signature. The connectivity score aims to summarize the transcriptomic relationship between each compound and disease, such that a strongly negative score indicates that the compound will induce transcriptomic changes that may revert or “normalize” the disease signature *(31)* Using this method, significant negative connectivity scores were found between the Src kinase inhibitor, saracatinib, and IPF disease signatures derived from patient lung biopsies (GSE24206) (34) and cultured fibroblasts (GSE44723) (35) (FDR <0.01), **(Table-S1)**.

Accordingly, we performed a disease enrichment analysis (DEA), with the goal of identifying high-level disease categories that are transcriptomically connected to saracatinib. This analysis identified that IPF disease signatures were highly over-represented in connecting with saracatinib (**Figure 1B**), indicating that saracatinib is globally relevant to IPF disease.

**Figure 1.**
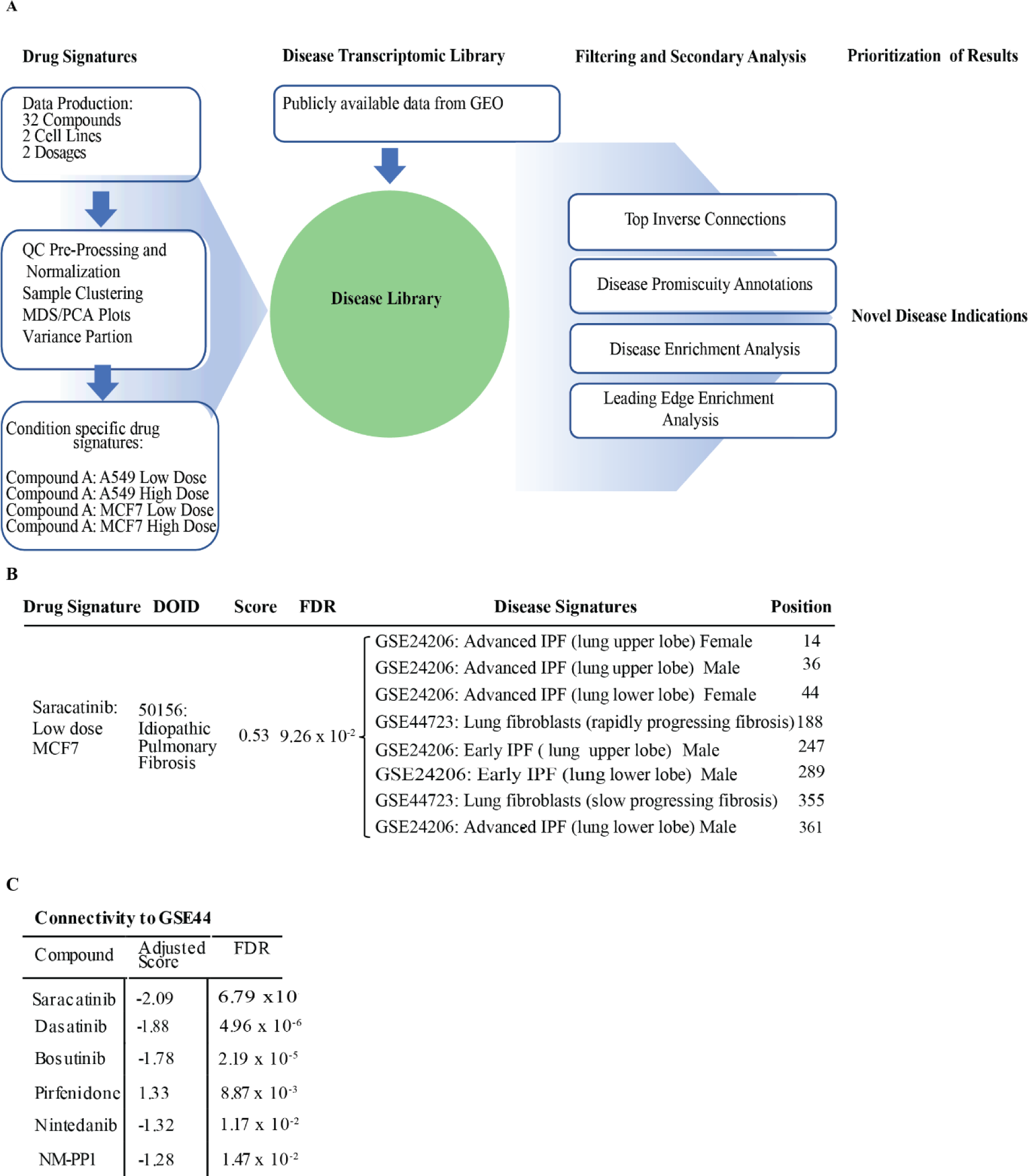
Computational drug repurposing identifies saracatinib as a potential therapeutic for IPF. **(A)** Schematic of the *in-silico* approach used to identify novel disease indications for compounds. Drug signatures were obtained for 32 compounds in two different cell lines at two dosages. Each drug signature was compared to a library of disease signatures generated from publicly available data and a connectivity score was generated for each disease-compound pair. Filtering and secondary analyses were carried out to identify novel disease indications for each of the compounds. **(B)** Disease enrichment analysis results showing enrichment of DOID:50156/IPF signatures among disease signatures that are transcriptomically connected to saracatinib. **(C)** Connectivity scores between an IPF disease signature and publicly available drug signatures (saracatinib, dasatinib, bosutinib, pirfenidone, nintedanib and NM-PP1; obtained from LINCS L1000).

In order to probe the biology driving the connection between saracatinib and the IPF signatures, we conducted a leading-edge enrichment analysis to identify over-represented gene sets that point to pathways by which saracatinib may affect IPF disease. This analysis identified numerous gene sets including interferon gamma (IFNγ) response, epithelial-mesenchymal transition (EMT), and tumor necrosis factor alpha (TNFα) signaling pathways, all of which have been implicated in the pathogenesis of IPF. Additionally, Kinase Enrichment Analysis (KEA) of these data identified enrichments for receptor-interacting serine/threonine-protein kinase 3 (RIPK3) along with multiple members of the mitogen activated protein (MAP) kinase family (Table 1).

**Table 1.** Leading edge enrichment analysis. Enrichment analysis performed on leading edge of saracatinib drug signature versus IPF disease signature (GSE24206). The drug-disease pair which showed the highest connectivity was used for the analysis, namely “Saracatinib MCF7 Low Dose” versus “Advanced_IPF_explant_upper_lobe obtained from GSE24206”. F.C.=Fold Change. Gene sets and overlapping genes are listed.

| Enrichments: ↑ IPF, ↓ Saracatinib |  |  |  |  | Enrichments: ↓ IPF, ↑ Saracatinib |  |  |  |  |
| --- | --- | --- | --- | --- | --- | --- | --- | --- | --- |
| Libra ry | Gene Set | FDR | Genes in Overlap | F.C. | Gene Set | FDR | Genes in Overlap | F.C. |  |
| Hallm ark | IFNγ response | 1.79 x 10<sup>-12</sup> | ADAR, CD74, HLA-A, HLA-B, HLA-DMA, HLA-G, ICAM1, IFI30, IFIT3, IFITM3, IRF1, LGALS3BP, LY6E, MT2A, MX1, MYD88, NFKBIA, PTPN6, SERPING1 | 9.48 | Myc targets | 3.47 x 10<sup>-9</sup> | CCT2, DEK, DHX15, ERH, GLO1, HNRNPA1, IARS, LDHA, NPM1, PSMA6, RPL6, RPLP0, SLC25A3, YWHAQ | 9.28 | |
|  | EMT | 5.94 x 10<sup>-11</sup> | CAPG, COL1A1, COL4A2, COL6A2, CYR61, FLNA, FN1, GEM, HTRA1, ID2, IGFBP2, IGFBP4, SERPINE1, SERPINH1, TAGLN, TGFBI, VIM | 8.54 | Allograft rejection | 1.35 x 10<sup>-4</sup> | B2M, HIF1A, IFNGR1, MRPL3, NPM1, RPS3A, TIMP1 | 8.59 |  |
| | TNFα signaling via NF-κB | 4.03 x 10<sup>-10</sup> | CCND1, CD83, CYR61, EGR1, FOSB, GEM, ICAM1, ID2, IER2, IRF1, JUNB, KDM6B, LAMB3, LIF, NFKBIA, SERPINE1, SLC2A3, ZFP36 | 7.88 | Apoptosis | 4.39 x 10<sup>-4</sup> | CAV1, GPX3, IFNGR1, PPT1, TGFB3, TIMP1, TXNIP | 6.91 | |
|  | Comple ment | 2.50 x 10<sup>-8</sup> | APOC1, C3, COL4A2, CSRP1, CTSD, CTSH, DUSP6, FN1, GNAI2, GNB2, IRF1, PFN1, SERPINE1, SERPING1, TIMP2 | 7.81 | Inflammator y response | 1.49 x 10<sup>-2</sup> | ADM, HIF1A, LAMP3, NAMPT, TIMP1 | 6.08 |  |
|  | Coagulat ion | 3.70 x 10<sup>-8</sup> | APOC1, C3, CRIP2, CSRP1, CTSH, DUSP6, FN1, GNB2, HTRA1, SERPINE1, SERPING1, USP11 | 10.07 | Reactive oxygen species | 1.81 x 10<sup>-2</sup> | GPX3, PRDX1, PRDX6 | 8.95 |  |
| | INFα response | 1.61 x 10<sup>-6</sup> | ADAR, CD74, HLA-C, IFI30, IFIT3, IFITM3, IRF1, LGALS3BP, LY6E, MX1 | 9.03 | Oxidative phosphorylat ion | 1.84 x 10<sup>-2</sup> | COX6C, COX7A2, LDHA, NDUFA4, SLC25A3, SLC25A5 | 3.98 | |
|  | Allograft rejection | 2.79 x 10<sup>-5</sup> | CAPG, CD74, FLNA, HLA-A, HLA-DMA, HLA-E, HLA-G, ICAM1, LIF, PTPN6 | 6.67 | Androgen response | 3.27 x 10<sup>-2</sup> | B2M, MYL12A, SGK1, SLC38A2 | 5.77 |  |
|  | Apical junction | 4.75 x 10<sup>-5</sup> | ACTB, ACTG1, ACTN4, CAP1, EVL, GNAI2, ICAM1, LAMB3, PFN1, TGFBI, VASP, ZYX | 5.63 | EMT | 4.94 x 10<sup>-2</sup> | CD59, GJA1, TGFB3, TIMP1 | 3.70 |  |
| KEA | RIPK3 | 3.42 x 10<sup>-6</sup> | ACTB, ACTG1, ACTN4, FLNA, GNB2, HSPA8, RPS27, AUB52, UBB, VIM | 8.31 | MAP3K14 | 3.89 x 10<sup>-10</sup> | HNRNPA1, HSPA1A, IARS, NPM1, RPL24, RPL4, RPL6, RPL7A, RPL10, RPS11, RPS23, RPS3A, RPS7 | 12.74 |  |
|  | MAP3K3 | 5.42 x 10<sup>-6</sup> | ACTB, ACTG1, ACTN4, FLNA, GNAI2, GNB2, HSPA8, NFKBIA, RPS27, TUBA1C, UBA52, UBB, VIM | 5.66 | RIPK3 | 3.42 x 10<sup>-6</sup> | CCT2, HNRNPA1, HSPA1A, PPP1CB, PPP1CC, PRDX1, SLC25A3, SLC25A5 | 12.17 |  |
|  | MAP3K1 | 1.56 x 10<sup>-5</sup> | ACTB, ACTG1, FLNA, HSPA8, LGALS3BP, NFKBIA, RPL35, RPS27, UBA52, UBB, VIM | 6.52 | MAP3K1 | 1.71 x 10<sup>-3</sup> | HSPA1A, RPL24, RPL4, RPS11, RPS23, SLC25A5 | 6.50 |  |
|  | MAP3K14 | 4.89 x 10<sup>-4</sup> | ACTB, ACTG1, FAU, HSPA8, NFKBIA, RPL35, RPS10, RPS18, RPS20 | 4.82 | MAP3K3 | 7.38 x 10<sup>-3</sup> | CCT2, HSPA1A, IARS, PPP1CC, SLC25A5, YWHAQ | 4.79 |  |

To clarify if this connectivity was a feature among other Src inhibitors or unique to saracatinib, we used data from the previously published connectivity Map, a collection of publicly available expression data from cultured human cells treated with small molecules *(32)* This data collection contains transcriptomic data in a range of experimental conditions for saracatinib as well as other Src kinase inhibitors. This resource also includes data from the two IPF FDA-approved drugs pirfenidone and nintedanib. We compared each of these drug signatures to the IPF disease signatures in our disease library. One of the IPF disease signatures connected significantly with all 6 compounds (saracatinib, dasatinib, bosutinib, pirfenidone, nintedanib and NM-PP1) and the strongest connection was with saracatinib (**Figure 1C**). In summary, using complementary bioinformatics approaches, we identified a robust transcriptomic connection between saracatinib and IPF thus providing a strong foundation for the hypothesis that saracatinib might have a potential therapeutic benefit in IPF.

### Saracatinib inhibits TGF-β–induced phenotypic changes in human lung fibroblasts, (In vitro)

As the compound signature was derived from initial cell screening with a computational approach, we next investigated the effects of saracatinib in a more disease-relevant setting to confirm the connection between saracatinib and IPF. We assessed the effect of saracatinib on TGF-β-induced fibrogenic processes in cultured primary normal human lung fibroblasts (NHLF) to study signaling pathways relevant to human IPF disease. We confirmed that, TGF-β stimulation induces a significant increase in Src phosphorylation of Y416, a response that correlates with activation of Src kinase activity*(36)* **(Figure S1A and S1B).** It has been well demonstrated that saracatinib treatment efficiently inhibits TGF-β–induced Src kinase activity in these cells*(16)*. We chose to compare the effect of saracatinib to the two FDA-approved anti-fibrotic drugs, nintedanib and pirfenidone in this *in vitro* system. We carefully selected the optimum dose for all three compounds based on the established clinically relevant doses*(37, 38)* and our initial screening experiments **(Figure S2A and S2B)**. We observed that saracatinib significantly inhibited TGF-β-induced expression of many profibrotic genes including *ACTA2*, *COL1A1* and *SERPIN1* to a similar or greater extent than that observed for nintedanib or pirfenidone **(Figure 2A, 2B and 2C**). To validate this finding, we repeated the experiment in primary human lung fibroblasts isolated from three different donors and confirmed the consistency of these findings across all three donors **(Figure S3A and S3B)**. We next compared the effects of the three drugs on TGF-β-induced Smad3 phosphorylation in NHLF, as a readout of canonical TGF-β fibrogenic signaling. Saracatinib, but not nintedanib nor pirfenidone, significantly inhibited TGF-β-induced Smad3 phosphorylation (**Figure 2D**). We extended our investigation by comparing the effects of saracatinib to the other two drugs on TGF-β-induced morphological changes in human fibroblasts. Confocal immunofluorescence imaging of these cells for alpha smooth muscle actin (α-SMA) and filamentous actin (F-actin) demonstrated a strong inhibitory effect of saracatinib on TGF-β-induced alterations in cell shape and stress fiber formation characteristic of myofibroblast transformation (fold change >2, p-value ≤0.001), **(representative images,** **Figure 2E**) and **(quantifications, Figure 2F and 2G)**. Nintedanib exhibited a similar, but less potent, inhibitory response compared to saracatinib on these TGF-β induced phenotypic changes. Concordant with our earlier findings, pirfenidone did not show any effects on TGF-β induced structural changes in human lung fibroblasts.

**Figure 2.**
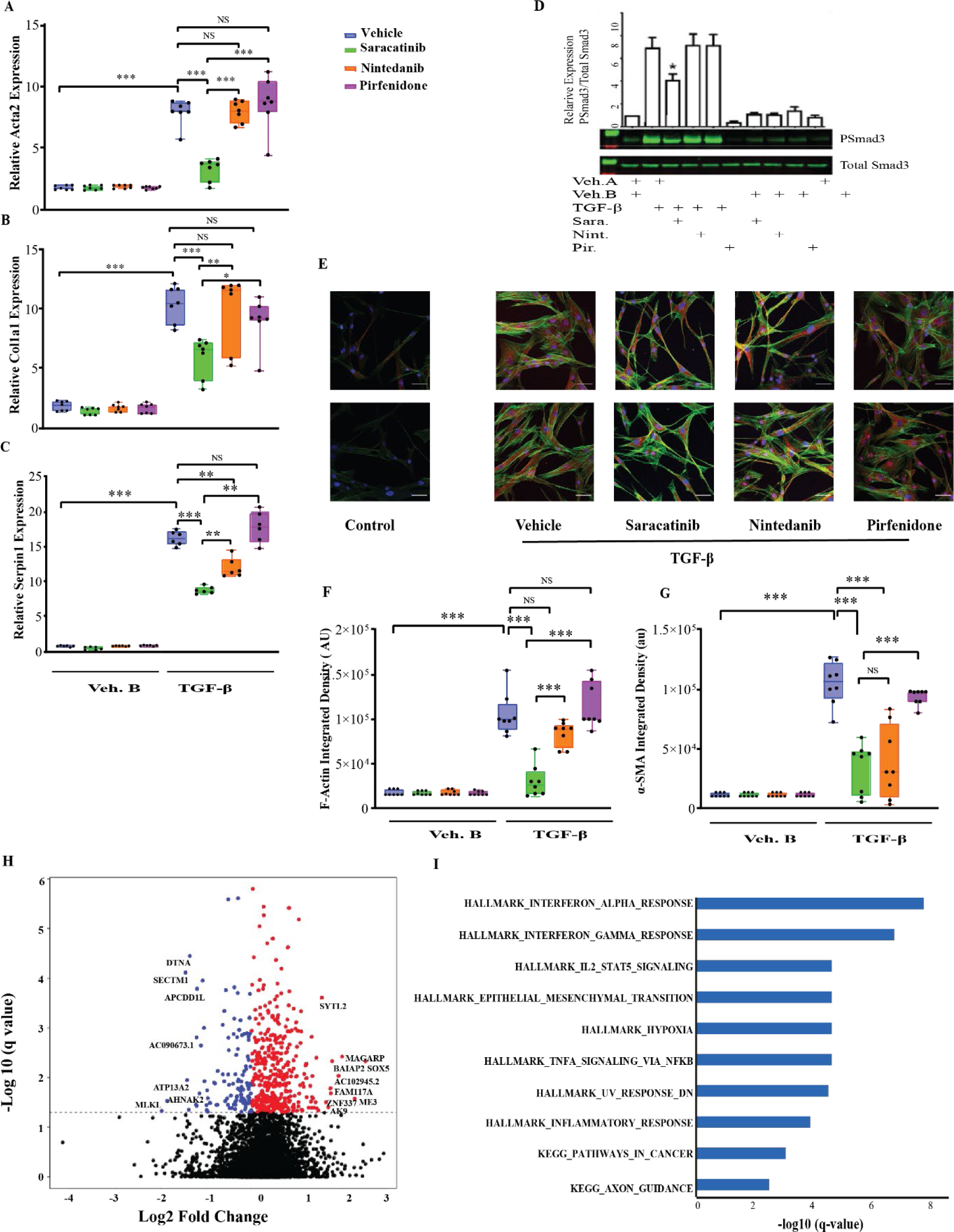
Saracatinib inhibits TGF-β–induced phenotypic changes in human lung fibroblasts (NHLFs). Cells were serum-starved overnight and then incubated with inhibitors (saracatinib: (0.3 μM), nintedanib: (1 μM), pirfenidone: (20 μg/ml) or Vehicle (DMSO)) for 60 min followed by stimulation with human recombinant TGF-β (2 ng/ml) or vehicle control for the indicated times.**(A, B and C)** RT-qPCR analysis for **(A)** *ACTA2* **(B)** *COL1A1* and **(C)** *SERPIN-1 (Pie-1)*, in the indicated treatment groups of NHLF(means+ SEM), *P<0.05,**P<0.01,***P<0.001(n=6).**(D)** Representative western blots showing saracatinib inhibits TGF-β-Induced phosphorylation of Smad3 in human lung fibroblasts, data presented as means+ SEM, n=6, *P<0.05 **(E)** Representative images of α-SMA staining (red) along with F-Actin (green) and DAPI (blue) show fluorescent staining in human fibroblasts after TGF-β stimulation in the indicated treatment groups using confocal microscopy. **(F)** and **(G)** Quantification of α-SMA and F-Actin staining shown as integrated density, ***P<0.001. **(H)** Volcano plot showing genes that are differentially expressed in cells treated with TGF-β and saracatinib compared to TGF-β alone (FDR<0.05, negative fold change (blue) and positive fold change (red). **(I)** Functional enrichment of significantly differentially expressed genes (FDR<0.05) in response to saracatinib (only top 10 gene sets shown. All gene sets shown are significant at FDR <0.05 and are from Hallmark (H) or Kegg (K).

To extend these observations and investigate the broader effects of saracatinib in this disease-relevant *in vitro* model, we generated transcriptomic drug signatures by performing bulk RNA sequencing (RNAseq) on NHLF cells treated with saracatinib, nintedanib, pirfenidone, or vehicle control in the presence or absence of TGF-β stimulation. We identified that saracatinib altered expression of over 500 individual genes (adj. p-value <0.05) in TGF-β treated cells (**Figure 2H**). We carried out gene set enrichment analysis (GSEA) with the goal of identifying gene sets that were significantly over-represented. This analysis revealed that saracatinib induced alterations in numerous pathways identified from the Hallmark and KEGG databases, with IFNα, IFNγ, EMT, and inflammatory responses among the top pathways (**Figure 2I** and **Table S2)**. We next identified the transcriptomic effects that were unique to saracatinib and could be used to differentiate the effects of saracatinib from nintedanib and pirfenidone **(Figure S4A and S4B)**. GSEA suggested that both nintedanib and saracatinib target several common gene sets including IFNγ, IFNα, and inflammatory responses. Gene sets that were uniquely enriched by saracatinib included EMT, TGF-β and WNT signaling while gene sets that were uniquely enriched by nintedanib related predominantly to metabolism. In summary, saracatinib inhibits TGF-β–induced fibrogenic responses in this *in vitro* system more effectively than nintedanib and pirfenidone.

### Saracatinib inhibits pulmonary fibrosis in preclinical animal models, (In vivo)

Next, we compared the effects of saracatinib, nintedanib and pirfenidone at clinically relevant doses *(39-43)*, in two preclinical models of pulmonary fibrosis in mice *(44, 45)*. In the first model, fibrosis was induced using a single dose of bleomycin (1.5U/Kg) administered into the lung by oropharyngeal aspiration (OPA) and mice received either saracatinib, nintedanib, pirfenidone or vehicle control once daily via oral gavages on days 10-27 post-bleomycin administration. On day 28, the mice were euthanized, lungs were harvested and the anti-fibrotic effects of all three drugs were assessed. Mice receiving vehicle control exhibited prominent weight loss as expected after the administration of bleomycin, while all drug-treated mice, except the mice treated with pirfenidone, recovered their weight loss by the end of the experiment **(Figure S5A and S5B)**. The bleomycin-induced increase in lung collagen content was significantly attenuated in mice receiving saracatinib, nintedanib, or pirfenidone (p-value ≤0.001) (**Figure 3A**). Consistent with the biochemical analysis, saracatinib significantly attenuated the bleomycin-induced whole lung expression of fibrogenic genes including *Acta2, Col1a1*, and *Col3a1* **(Figure 3B, 3C** and **S6A)**, (p-value ≤0.001 for *Acta* 2 and *Col3a1*, p-value ≤0.01 for *Col1a1)*, whereas neither nintedanib nor pirfenidone treatment resulted in any significant changes in the expression of these fibrogenic genes. Histopathological evaluation of these lungs using Masson’s Trichrome staining demonstrated a greater reduction in bleomycin-induced pulmonary fibrosis by saracatinib and nintedanib (p-value ≤0.001) compared to pirfenidone (p-value ≤0.01) **(representative images,** **Figure 3D** and **quantifications,** **Figure 3E**). Micro-computerized tomography (CT) analysis of these mice lungs supported the biochemical and histopathological observations; aerated lung volume measurements showed saracatinib and nintedanib significantly attenuated the bleomycin-induced radiographic alterations in lung parenchyma (p-value ≤0.001) and to a greater extent than pirfenidone (p-value ≤0.01) **(representative images,** **Figures 3F, S7** and **quantifications, Figure, 3G)**. Finally, lung function measurements and physiological evaluation of these mouse lungs identified that saracatinib also significantly attenuated bleomycin-induced alterations in lung physiology [static compliance (Cst) and elastance] as measured using the flexiVent system (fold change=2, p-value ≤0.001) compared to nintedanib or pirfenidone (p-value ≤0.01) (**Figure 3H**).

**Figure 3.**
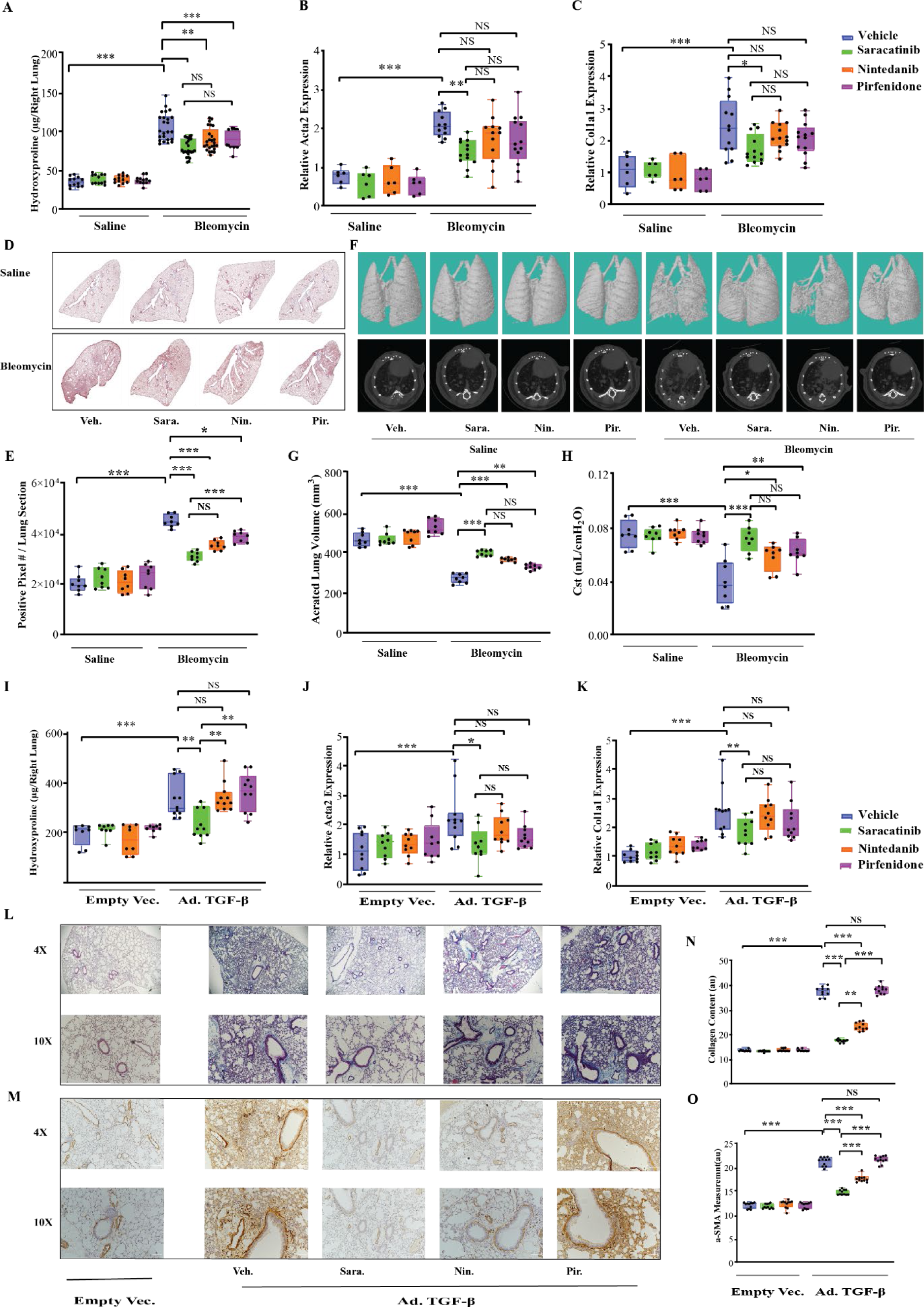
Saracatinib inhibits pulmonary fibrosis in bleomycin and adenovirus TGF-β mouse models. **(A-H) Evaluation of bleomycin-induced lung fibrosis: (A**) Quantitative analysis of hydroxyproline in lung homogenates from indicated groups of mice. Lung collagen content increased significantly in bleomycin-treated mice receiving vehicle control (fold change=2.9), **P < 0.01, ***P < 0.001. **(B and C)** RT-qPCR analysis on mouse lungs for **(B)** *Acta2* **(C)** *Col1a1* in the indicated treatment groups. **(D and E)** Representative images and quantitative measurements of Masson’s Trichrome staining of lung sections in the indicated groups of mice. **(F and G)** Representative images (dorsal view of three -dimensional reconstructions and axial view) and quantifications of microCT on mouse lung tissues in the indicated groups. Gross abnormality resulting from bleomycin-induced lung fibrosis is alleviated after treatment. Aerated lung volume measurements showed a significant decrease in bleomycin-treated mice lung (p-value ≤0.001, fold change >2), while saracatinib and nintedanib significantly attenuated the bleomycin-induced radiographic alterations in lung parenchyma (p-value ≤0.001). **(H)** Lung compliance measurements of each mice lung at the end of the study in the indicated groups, shown as static compliance (Cst). **(I-O) Evaluation of Ad-TGF-β-induced lung fibrosis: (I)** Quantitative analysis of hydroxyproline in lung homogenates from indicated groups of mice, the hydroxyproline assay revealed a significant increase in lung collagen content for mice receiving Ad-TGF-β (fold change=1.8, p-value ≤0.001) **(J and K)** RT-qPCR analysis on mouse lungs for **(J)** *Acta2* **(K)** *Col1a1* in the indicated treatment groups. **(L and N)** Representative images and quantification of Masson’s Trichrome staining of lung sections in the indicated groups of mice. **(M and O)** Representative images and quantification of α-SMA staining of lung sections in the indicated groups of mice. All data is presented as (means+ SEM), *P <0.05, **P <0.01, ***P<0.001, (n=6 in controls and n≥12 in bleomycin and TGF-β treated groups).

A second model of pulmonary fibrosis was established using expression of recombinant murine TGF-β using adenoviral-mediated gene delivery (Ad-TGF-β) administered to the lung intranasally. As in the bleomycin model, all the three drugs were given to mice once daily via oral gavages from days 10-27 after TGF-β administration. Similar to the bleomycin model, mice receiving vehicle control exhibited prominent weight loss after the administration of TGF-β adenovirus, but notably, only the mice treated with saracatinib recovered their weight loss by the end of the experiment (Figure S3C and S3D). In this model, only saracatinib demonstrated antifibrotic effects as assessed by a reduction in lung collagen content (p-value≤ 0.001) (**Figure 3I**) and expression of *Acta2*, *Col1a1* and *Col3a1* mRNA (p-value ≤ 0.001) (**Figure 3J** and **3K** and S6B). By contrast, neither nintedanib nor pirfenidone attenuated fibrosis in this model. Histopathological assessment of the lungs using Masson’s Trichrome and α-SMA staining confirmed these observations (representative images, **Figure 3L, 3M**, S8A and S8B and quantification, **Figure 3N** and **3O**).

Given the fact that TGF-β signaling cascades are consistently activated in fibrotic tissues, regardless of the etiology of the initial injury and that Smad3 pathway is a key intermediary in this cascade *(46, 47)*, we evaluated the effect of saracatinib on Smad3 phosphorylation in the lung tissues after bleomycin and Ad-TGF-β treatment. We identified significant inhibition of this pathway by saracatinib in both preclinical animal models of pulmonary fibrosis (Figure S9A-D).

In summary, in two complementary preclinical mouse models of IPF, saracatinib attenuated experimental pulmonary fibrosis significantly more effectively than pirfenidone or nintedanib at clinically relevant doses.

### Saracatinib inhibits lung fibrosis in precision cut lung slices (ex vivo model)

We next sought to confirm these observations in an *ex vivo* model using precision cut lung slices (PCLS) to compare the anti-fibrotic effects of saracatinib with nintedanib and pirfenidone *(48)*. PCLS is a useful method to test the therapeutic effects of different compounds in a relatively preserved lung architecture containing various lung-resident cell types and ECM *(49)*. In this study, mice were treated with either bleomycin or Ad-TGF-β and euthanized at the peak of the fibrotic response (day 14 for bleomycin and day 21 for Ad-TGF-β model). Lung slices were prepared and treated *ex vivo* with each of the three drugs or vehicle for 5 days (**Figure 4A-L**). These lung slices remain viable at least for 5 days (120 hours), **(Figure S10)**. Quantitative RT-PCR (qRT-PCR) analysis of lung slices revealed a dramatic reduction in *Col1a1* and *Acta2* mRNA expression by saracatinib in PCLS harvested from mice treated with either bleomycin or Ad-TGF-β (p-values≤0.001), while the fibrosis was sustained in the control groups (**Figure 4A** and **4B; 4G** and **4H,** respectively). Live serial imaging of the lung slices using second harmonic generation microscopy (SHG) demonstrated that saracatinib significantly attenuated collagen accumulation in both models (p-values≤0.001) **(representative images, Figure 4C** and **4I** and **quantifications, Figure 4D** and **4J)**. Note that it was not possible to assess the effects of nintedanib on collagen content by SHG in this model due to a strong auto-fluorescence signal generated by nintedanib. Treatment with pirfenidone attenuated lung collagen content (p-values≤0.001) only in the TGF-β model. Histological evaluation of the lung slices using Masson’s Trichrome staining revealed that saracatinib strongly attenuated lung fibrosis in both models (p-values≤0.001) **(representative images, Figure 4E and 4K** and **quantifications, Figure 4F** and **4L)**. In summary, using PCLS as a complementary model, saracatinib was more effective than either nintedanib or pirfenidone in attenuating pulmonary fibrosis in an *ex vivo* model and confirmed our finding in the *in vivo* murine models.

**Figure 4.**
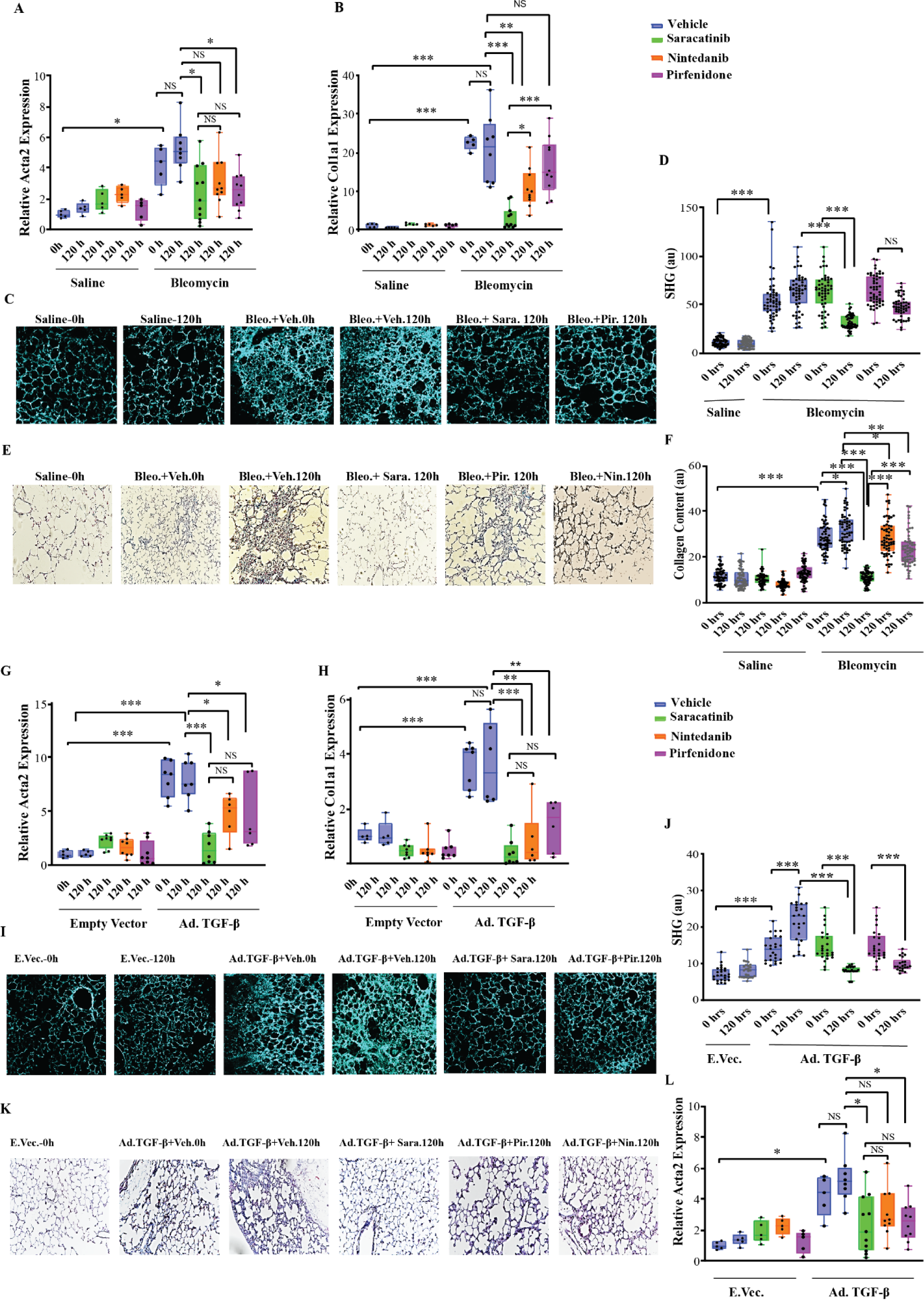
Saracatinib inhibits pulmonary fibrosis in *ex vivo* murine PCLS in bleomycin and Ad-TGF-β models. Mouse precision cut lung slices (PCLS) were generated from both bleomycin (day 14) and Ad-TGF-β (day 21). Treatment with saracatinib (0.6μM), nintedanib (1μM), pirfenidone (1mM) or vehicle control, was administered in the first 24 hours after slicing (Time 0 hour). All lung slices were isolated 5 days after treatment for analysis (Time 120 hours). **(A-F)** Effects of saracatinib, nintedanib and pirfenidone in PCLS isolated from bleomycin mice model of pulmonary fibrosis. **(A and B)** RT-qPCR analysis on mice PCLS in bleomycin model for *Acta2* and *Col1a1* at time 0 and 120 hours after indicated treatments. **(C and D)** Representative live images by second harmonic generation microscopy (SHG) and quantification assessments of PCLS samples at time 0 and 5 days (120 hours) after indicated treatments. **(E and F)** Representative images and quantification assessments of Masson’s Trichrome staining of PCLS slides at time 0 and 5 days (120 hours) after indicated treatments. **(G-L)** Effects of saracatinib, nintedanib and pirfenidone in PCLS isolated from Ad-TGF-β treated murine model of pulmonary fibrosis. **(G and H)** qRT-PCR analysis on murine PCLS in the Ad-TGF-β model for *Acta2* and *Col1a1* at time 0 and 120 hours after indicated treatments. **(I and J).** Representative live images using second harmonic generation microscopy (SHG) and quantification assessments of PCLS samples at time 0 and 5 days (120 hours) after indicated treatment. **(K and L)** Representative images and quantification assessments of Masson’s Trichrome staining of PCLS samples at time 0 and 5 days (120 hours) after indicated treatment. All data is presented as (means+ SEM), P < 0.05, **P<0.01, ***P<0.001, (n≥6 in all groups).

### Comparison of transcriptional changes in mouse models identifies core genes relevant for IPF and points to a mechanism of action of saracatinib

We undertook bioinformatic analysis to explore the transcriptional changes underlying the observed anti-fibrotic effects of saracatinib in both murine models of pulmonary fibrosis. RNAseq was carried out on lungs isolated from mice treated with either bleomycin or Ad-TGF-β in the presence or absence of saracatinib treatment. Consistent with the favorable clinical safety profile of saracatinib, there were relatively few gene expression changes in mice receiving saracatinib alone compared to the control group. Importantly, saracatinib reversed the expression of many of the genes that were altered by bleomycin **(Figure 5A and 5B)**. This was also reflected by pathway analysis; GSEA demonstrated that saracatinib reversed alterations in numerous gene sets induced by bleomycin (including Myc Targets, E2F Targets, EMT, and G2M Checkpoint gene sets) (**Figure 5C**). Analysis of the pulmonary transcriptional changes observed in the Ad-TGF-β model revealed that TGF-β induced changes in approximately 4500 genes; however, contrary to the observed changes in the bleomycin model, saracatinib treatment had a much smaller effect with fewer significant changes in expression. Further evaluation using GSEA, taking expression levels of all genes into account, demonstrated that saracatinib attenuated many pathways altered by TGF-β expression including IFNγ, IL6, JAK-STAT3 signaling, and IFNα responses (**Figure 5D**).

**Figure 5.**
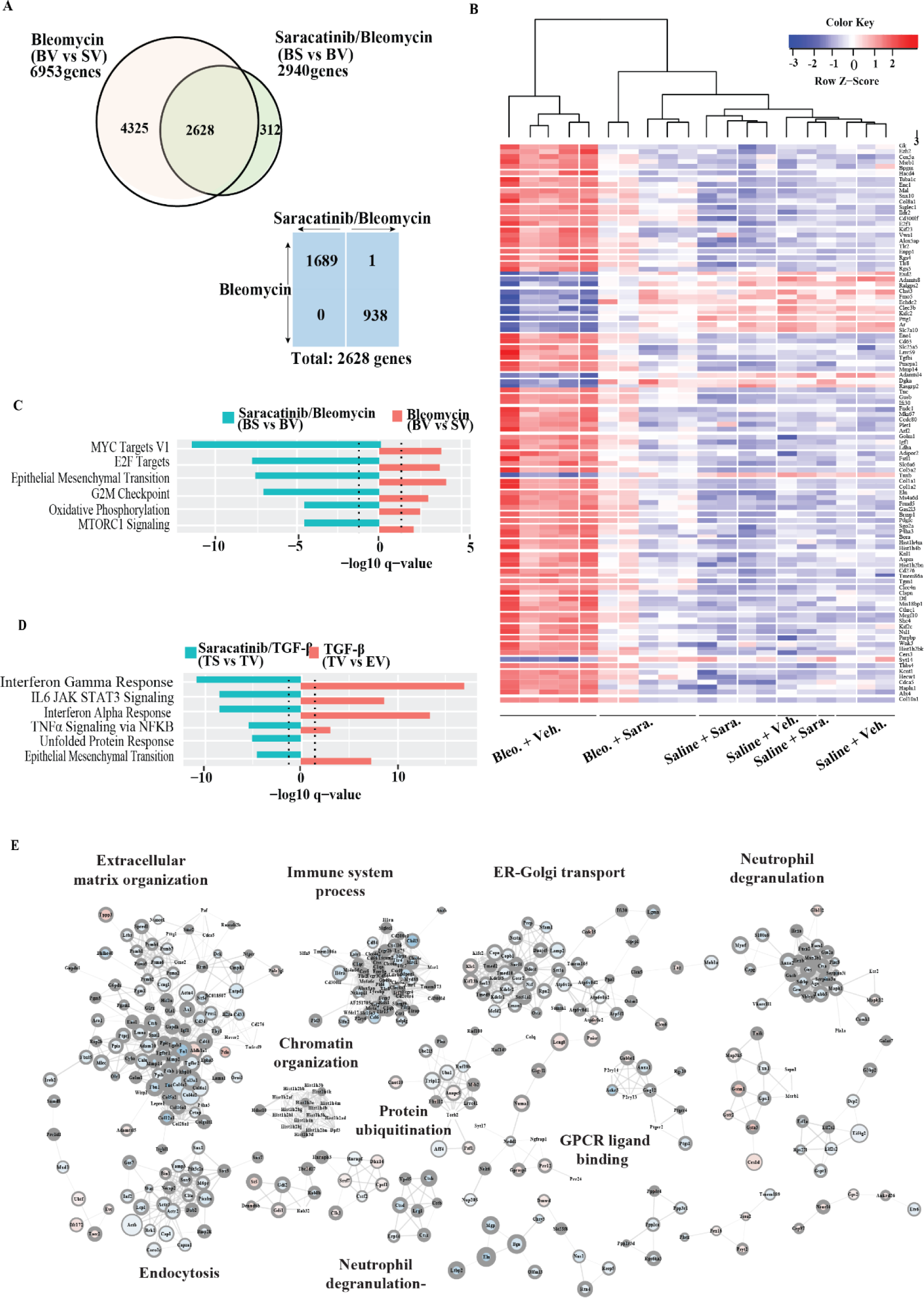
Saracatinib treatment results in reversal of transcriptional changes observed in IPF mouse models. **(A)** Comparison of number and direction of significantly differentially expressed genes (FDR <0.05) in bleomycin, and bleomycin/saracatinib treatment groups. Bleomycin administration induced significant differential expression of almost 7000 genes (adj. p-value <0.05) in the bleomycin treatment group when compared the control group. Bleomycin-treated mice that also received saracatinib had significant changes in 2940 genes compared to mice treated with bleomycin alone. 2628 differentially expressed genes are common between treatment groups. Of these 1689 are upregulated by bleomycin and downregulated by saracatinib; and 938 are downregulated by bleomycin and upregulated by saracatinib. (**B)** Heatmap of top 100 differentially expressed genes (BS vs BV). **(C)** Top Hallmark GSEA in bleomycin murine experiments (adj. p-value <0.05) ranked by saracatinib effect. Blue bars show BS vs BV and red bars show BV vs SV. The positive or negative signs indicate logFC directions. **(D**) Top Hallmark GSEA in Ad-TGF-β experiments ranked by saracatinib effect (adj. p-value <0.05). Blue bars show TS vs TV and red bars TV vs EV. The positive or negative signs indicate logFC directions. **(E)** Cytoscape network analysis of common DE genes shared by both the bleomycin and TGF-β mouse models that are reversed by saracatinib. Node size - expression level; Node color – LogFC (red is up, blue is down); node border width – negative log adj. p-value. *<u>Key: Bleomycin experimental conditions:</u> SV: saline + vehicle; BV: bleomycin + vehicle; BS - bleomycin + saracatinib; BV vs SV - bleomycin effect; BS vs BV - saracatinib on bleomycin. Ad-<u>TGF-β experimental conditions</u>: EV: control + vehicle; TV: TGF-β + vehicle; TS: TGF-β + saracatinib; TV vs EV: TGF-β effect; TS vs TV: saracatinib on TGF-β*.

As neither mouse model faithfully recapitulates all aspects of human IPF, we compared the transcriptional changes observed in these two animal models and identified a set of genes common to both models (consisting of 1233 upregulated genes and 1256 downregulated genes). A subset of these genes is reversed by saracatinib (225 upregulated, 560 downregulated). To interrogate this subset further, a protein-protein interaction network analysis identified modules including ECM organization, immune system processes, ER-golgi transport and neutrophil degranulation (**Figure S10**). Within this ECM cluster are numerous known markers of fibrosis including members of the collagen family *(Col3a1, Col4a1, Col4a2, Col5a2, Col12a1, Col16a1)*, fibronectin *(Fn1)*, laminin *(Lama1)*, tenascin *(Tnc)*, fibrillin *(Fbn1)*, TGF-β receptor1 *(TGF-βr1)* and the ECM receptor *Cd44*(Figure S11).

## Discussion

In this study, we used an innovative disease-agnostic computational biology-based approach and identified the Src kinase inhibitor, saracatinib (AZD0530) *(29)*, as a potential therapeutic agent for IPF and delineated multiple fibrogenic pathways targeted by this agent at the molecular level. Further, we validated the efficacy of saracatinib in blocking fibrogenic responses in several complementary pre-clinical models of pulmonary fibrosis. Our data provide strong evidence that saracatinib is equal or superior to the two FDA-approved drugs, nintedanib and pirfenidone at inhibiting pulmonary fibrosis in experimental models. Analysis of the transcriptional changes induced by saracatinib in the *in vitro* and *in vivo* models of pulmonary fibrosis demonstrated that saracatinib was able to reverse the expression of diverse fibrogenic genes and pathways such as myofibroblast differentiation, EMT, ECM organization, immune system processes, ER-golgi transport, and neutrophil degranulation. Importantly, many of these differentially expressed gene sets and pathways are selectively modified by saracatinib as compared to nintedanib and pirfenidone. Taken together, our data support the validity of a transcriptomics-based bioinformatics approach to generate novel hypotheses for drug repurposing in complex human diseases such as IPF.

At the inception of this study, we used genome-wide transcriptional profiling of human IPF lungs cross-referenced with distinct drug transcriptomic signatures to identify Src kinase-dependent signaling pathways as critical checkpoints that control pro-fibrotic responses in the lung. Importantly, we established that saracatinib was a potent and effective therapeutic agent that significantly attenuated experimental pulmonary fibrosis in several preclinical models with equal or greater efficacy compared to the two FDA-approved drugs, nintedanib and pirfenidone. This approach, anchored in transcriptomic profiles in human lung cells and tissues, led to identification of novel pathways that are relevant to human IPF.

Our study provides additional information and is complementary to proteomic-based analyses of human tissues that have yielded therapeutic targets for pulmonary fibrosis *(50)* This is of considerable practical importance because, despite massive investments made by academia and industry, many drugs have failed in later stages of development, partly due to the poor predictability of animal models in drug discovery that do not reflect human disease *(51,52)*.

Repurposing drugs to treat conditions other than the one for which they were developed and/or approved, can shorten timelines, decrease costs, and increase success rates. Using a computational biology approach in combination with Connectivity Map (CMap) analysis will facilitate identification of new drugs, predicting drug candidates, and discovering connections among small molecules sharing a mechanism of action. Connectivity MAP analysis of single cell RNA sequencing of bronchial brushings from IPF patients also suggested that gene expression changes in IPF airway basal cells can be reversed by SRC inhibition, (https://www.biorxiv.org/content/10.1101/2020.09.04.283408v1). Based on the connectivity scores of saracatinib to IPF relative to nintedanib and pirfenidone using L1000 data, we identified that saracatinib had greater connectivity to IPF than the other two FDA approved drugs. This approach has been widely used as a resource in cancer drug discovery *(31, 53)* and our study extends this methodology to IPF treatment.

Several studies have shown that SFK control key signaling pathways relevant to pulmonary fibrosis pathogenesis *(54)* including TGF-β-mediated myofibroblast differentiation and fibrogenic responses in lung fibroblasts *(16)*. SFK are also required for TGF-β-driven EMT and mediate TGF-β-induced β-catenin phosphorylation, a critical determinant of fibrogenesis in alveolar epithelial cells *(55)*. SFK are also necessary for the recruitment and activation of immune cells as well as having other roles in pulmonary inflammation *(54)* We have reported that mice genetically deficient in protein tyrosine phosphatase (PTP)α, a known activator of SFK, are protected from experimental pulmonary fibrosis *(56, 57)*. We also reported that the SH2 domain–containing tyrosine phosphatase2 (SHP2) is an antifibrotic regulator in pulmonary fibrosis *(58,59)*. These studies underscore the importance of reversible tyrosine phosphorylation reactions, controlled by specific tyrosine kinases and phosphatases, in the pathogenesis of pulmonary fibrosis and provide a mechanistic foundation for the use of saracatinib in the treatment of pulmonary fibrosis.

Our studies using normal human lung fibroblasts provide strong evidence that saracatinib inhibits TGF-β-induced fibrogenic responses, more potently than either nintedanib or pirfenidone, at the levels of mRNA expression, post-translational signal transduction, and myofibroblast morphological changes. We observed that saracatinib was more effective than nintedanib or pirfenidone in animal models and in PCLS. Both models revealed that saracatinib perturbed ‘EMT’, ‘MYC targets’ and ‘IL6_JAK_STAT signaling’ gene sets and that saracatinib uniquely induced perturbations in ‘E2F target’, ‘G2M checkpoints’ and ‘oxidative phosphorylation’ gene sets in the bleomycin model, and interferon gene sets in the Ad-TGF-β model. This analysis provides important insights into the mechanisms by which SFK modulate fibrogenic signaling pathways. Further investigation into the precise mechanisms of saracatinib action in the context of IPF is the subject of ongoing/future work.

Several questions of interest with respect to the effects of saracatinib on fibrogenic pathways remain to be addressed. In the current study we focused on TGF-β-dependent pathways because of the well accepted importance of this growth factor in organ fibrosis *(60,61)*. In addition to TGF-β, other growth factors including platelet-derived growth factor (PDGF) *(62,63)*, insulin-like growth factor-(IGF)1 *(64)* and fibroblast growth factors (FGFs) *(65)* are known to trigger pro-fibrotic cellular responses through signaling pathways that are controlled by SFK. In future studies, we plan to investigate the effects of saracatinib on PDGF, IGF-1, and FGF fibrogenic signaling.

Kinase Enrichment Analysis (KEA) of differentially expressed gene sets altered by saracatinib identified enrichment for RIPK3 and the MAP Kinases. Importantly, RIPK3 has been implicated in the pathogenesis of renal fibrosis *(66)* SFK are known to regulate MAP Kinase-dependent signaling pathways relevant to fibrogenesis *(67,68)*. Whether RIPK3 is regulated by SFK is unknown and the subject of current investigations.

In conclusion, based on a computational drug repurposing strategy, we have identified saracatinib as a potential therapeutic for IPF. As the safety profile of saracatinib has previously been established, the efficacy of saracatinib in IPF patients is now being investigated in a Phase 1b/2a clinical trial (NCT04598919). To our knowledge, this is the first study that has used a computational approach to link a compound to a specific disease using a computational approach, validated the efficacy of the drug in three complementary preclinical *in vitro*, *in vivo* and *ex vivo* models and culminated in a human clinical trial in the treatment of IPF.

## Materials and Methods

### Reagents

Saracatinib was provided by AstraZeneca. Pirfenidone and nintedanib were purchased from Tocris Bioscience (Cat No. 1093) and Selleckchem (Cat No. S1010), respectively. Hydroxypropyl methyl Cellulose (HPMC) was purchased from Sigma (Cat No. P8074).

### In vitro assays

Experiments were performed using primary normal human lung fibroblasts (NHLF) from three different donors (Lonza) and all experiments were performed using cells of passages 3-5. All experiments were performed with 6 technical replicates and repeated at least three times.

### TGF-β-induced Src activation

Normal Human Lung Fibroblasts were plated on rat tail collagen-coated non-tissue culture plastic, serum starved overnight, and then stimulated with human recombinant TGF-β (2 ng/ml) for 60 min at 37°C. Protein was solubilized in RIPA buffer and then subject to SDS-PAGE and immunoblotting with antibodies to total SRC and phospo-Y416 SRC (Cell Signaling, 2110 and 6943 respectively). Membranes were scanned using a Licor Odyssey fluorescent imaging instrument and densitometry conducted using Image Studio Lite, N=6, ** P=0.0055.

### Dose selection experiment

For choosing the optimal dose, cells were serum-starved overnight and then incubated with inhibitors at two selected clinically relevant doses (saracatinib: (0.12 & 0.3 μM), nintedanib: (0.1 &1 μM), pirfenidone: (10 and 20 μg/ml) or vehicle A (DMSO)) for 60 min followed by stimulation with human recombinant TGF-β1 (2 ng/ml) or vehicle B as control for 6 and 24 hours. After initial pilot experiments to identify optimal conditions, we selected clinically relevant concentrations for each drug [based on the known pharmacokinetic profile (Cmax) in humans available from the respective clinical development programs] and explored the effects of all three compounds on fibrogenic responses to TGF-β in normal human fibroblasts. Cells were serum-starved overnight and then incubated with inhibitors [saracatinib: 0.3 μM, nintedanib: 1 μM, pirfenidone: 20 μg/ml or vehicle (DMSO)] for 60 min followed by stimulation with human recombinant TGF-β (2 ng/ml) or vehicle control for the indicated times (30 min for western blots, 24 hours for gene expression) in the continued presence of the inhibitors.

### Gene Expression Analysis

For qPCR analysis, cells were lysed in Trizol, RNA was extracted with miRNeasy Mini Kit from Qiagen. The levels of mRNA for the fibrogenic genes were quantified by RT-qPCR and normalized to GAPDH expression.

### Western Blots

For western blot, cells were lysed in RIPA buffer and analyzed by SDS-PAGE followed by western blotting with antibodies to phospho and total Smad3 (Cell Signaling). The blots were quantified using an Odyssey fluorescent imaging system.

### Immunofluorescence (IF) staining

IF was performed 24 hours after the treatments, using anti-α-SMA (ab5694, Abcam) with T-6778 anti-rabbit Ig-TRITC as a secondary antibody, along with F-actin stain (Invitrogen-R37710), following the manufacturers’ protocols. Imaging was conducted by confocal microscopy (Leica SP5) and images were processed and quantified using Image J software.

### *In vivo* experiments

All animal procedures were approved by the Institutional Animal Care and Use Committees (IACUC) from National Jewish Health and Yale University. Pulmonary fibrosis was induced in C57Bl/6 mice by intrapulmonary delivery of bleomycin or recombinant adenovirus TGF-β (Ad-TGF-β). Bleomycin (1.5 U/kg) or saline administered by oropharyngeal instillation and Ad-TGF-β1 (VQAd CMV mTGF-β1-Viraquest), (2 x 10^9^ pfu per mouse) or empty vector was administered via the intranasal route. In both animal models, treatment with drugs was started on day 10 and drugs were administered daily by oral gavage (saracatinib 20mg/kg; nintedanib 60mg/kg; pirfenidone 300mg/kg) until day 28. Mice were euthanized, and lungs were harvested on day 28 for fibrosis analysis. Mice of both sexes were studied and 15-20 mice per group were used.

### Hydroxyproline Assay

Whole right lungs from each animal were used for hydroxyproline measurements using Hydroxyproline Colorimetric Assay Kit (Biovision K555-100). Briefly, constant weight homogenates from the mice lungs were hydrolyzed in 12 N HCl for 3 hrs at 120°C. The dried digestions reacted with Chloramine T and were visualized by DMAB reagents. The absorbance was measured at 560 nm in a microplate reader. Data were expressed as μg of hydroxyproline/right lung.

### Gene Expression

Left lungs were used for RNA extraction via miRNeasy Mini Kit (Qiagen-217004) according to the manufacturer’s instructions. The purity of the RNA was verified using a NanoDrop at 260 nm, and the quality of the RNA was assessed using the Agilent 2100 Bioanalyzer (Agilent Technologies). Purified RNA was used either for gene expression analysis by TaqMan gene expression assays or RNAseq analysis.

### Histology

Animal tissue sections from all the groups were stained with Masson’s Trichrome (collagen/connective tissue), H&E (Hematoxylin and Eosin stains) or α-SMA (alpha smooth muscle). Immunostaining was performed after paraffin removal, hydration, and blocking, following the recommendation of the manufacturer (ABC detection system from Vector lab, USA). All sections were analyzed using a Nikon microscope. Quantification of collagen was done with ImageJ software.

### MicroCT Analysis

MicroCT scan was performed on live animals using Bruker SkyScan 1276 Desktop *In-Vivo* MicroCT System following the manufacturers’ protocols. The microCT data from all the mice lungs were analyzed using the computer program CT-Analyser (CTAn) by Bruker. In brief, the CT scan is turned into a binary image and the threshold for what is considered aerated lung tissue vs other tissues is chosen by the blinded person performing the data analysis. This threshold is chosen based on closely matching the binary image of the scan to the actual scan. The same threshold is then applied to each scan. With CTAn, an automatic series of computer operations separates the aerated lung tissue from the rest of the mouse body, and this is quantitated in mm^3^. The Hounsfield Units are obtained with this same program from a calculation that is based on a CT scan of distilled water serving as a standard for the CT scanner. Reconstructed scans are corrected for misalignment, ring artifact, and beam hardening. The same beam hardening, and ring artifact corrections are applied to all scans. The misalignment is unique for each scan. This is done with NRecon from Bruker. The detailed explanation of aerated lung volume calculation includes a) Turn image into binary image and set ROI (region of interest) as the mouse body. b) Reload original image with new ROI as mouse body. c) Apply aerated lung vs everything else threshold to scan inside ROI. d) Perform a series of computer operations to remove everything but the aerated lungs. e) Data analysis on lungs, save lungs as ROI. f) Load lung only ROI. g) Obtain Hounsfield Unit (HU) measurement using correction factor obtained from Distilled (DI) water scan.

### Lung Function Test

Pulmonary function assessment was performed using Plethysmography and flexiVent (SCIREQ) systems following the manufacturers’ protocols on all animals at the end of the experiment.

### Western Blot

For Western blot, frozen tissues were lysed and homogenized in T-Per (Thermo Fisher Scientific) with phosphatase and protease inhibitor (100 µl per 10 mg tissue, abcam, ab201119). Protein content was measured with Nanodrop (280nm) and denaturation was performed at 95°C for 5 min in the presence of 2-mercaptoethanol and Laemmli buffer. 20 µg protein per lane was loaded onto a 4-20% gel (Bio-Rad) and samples were run at 50 mA followed by transfer on PVDF membranes using the Trans-Blot Turbo Transfer System (biorad). Membranes were washed, blocked in 5% dry milk (American bioInc) for 60 min followed by incubation overnight (at 4°C) with primary antibody according to manufacturer’s instruction (Cell Signaling Technology, pSMAD3 (9520S) SMAD3 (9513S), both 1:1000). Signal was detected using a donkey anti rabbit HRP conjugated antibody (Cytiva, 1:1000 for 1h at room temperature) using enhanced chemiluminescence substrate (biorad). Quantification was done using ImageLab software.

### *Ex vivo* Experiments, (Precision cut lung slices (PCLS))

Mouse precision cut lung slices were generated as previously described *(48, 69, 70)*. Mouse lungs from both bleomycin (day 14) and Ad-TGF-β (day 21) models were used in parallel and repeated accordingly. Briefly, low melting grade agarose (3 wt-%) was slowly injected via the trachea to artificially inflate the lung. Lungs were cooled at 4°C for 15 minutes to allow gelling of the agarose and then cut to a thickness of 150µm using a Compresstome (VF-300-0Z by Precisionary®) at cutting speed of 6 μm/s and oscillation frequency of 5Hz. The PCLS were cultured in a 24-well plate (Corning®) in 500μL DMEM-F12 no-phenol red containing 0.1% FBS and 1% penicillin/streptomycin 37 °C, 5% CO2 and 95% humidity. Treatment with saracatinib (0.6μM), nintedanib (1μM), pirfenidone (1mM) or vehicle control, was administered in the first 24 hours after slicing. All media were changed daily, and lung slices were isolated 5 days after treatment for analysis. Serial live imaging of lung slices by second harmonic generation microscopy technique was performed before (Time; 0 hours) and after each treatment (Time; 120 hours) *(71)* PCLS were fixed with 4% (weight/volume) paraformaldehyde overnight, and paraffin embedded at 0h and 120h. 3µm sections were cut using a microtome, mounted on glass slides and subjected to antigen retrieval. After deparaffinization and rehydration, staining was performed according to standard protocols for Masson’s Trichrome. Finally, samples were mounted using mounting medium and covered with a cover slip. Microscopic scanning of the slides was conducted in bright field with a Nikon inverted microscope at 20X magnification. 2 representative images were acquired for each sample and at least 20 different random field of views were used for collagen quantification. RNA was isolated from 5-6 replicated lung slices from each treatment using miRNeasy Mini Kit (Qiagen) at time 0 and time 120 hours for all groups. mRNA levels of fibrogenic genes were quantified by RT-qPCR and normalized to GAPDH expression. All experimental groups were performed in a group of 6 technical replicates and repeated at least three times.

### Live/Dead assay

Cell viability in PCLS was assessed using LIVE/DEAD® Viability/Cytotoxicity Kit following manufacturer’s instruction (ThermoFisher Scientific; Cat. No. L3224). PCLS were incubated with 2 µM Calcein AM and 2 µM Ethidium homodimer-1 (EthD-1) in 250 µm HBSS for 30 min at 37 °C. They were then washed two times with HBSS and fixed with 10% buffered formalin for 30 min at RT and washed. Samples were mounted on glass slides with ProLong® Gold Antifade Mountant (ThermoFisher Scientific, Cat. No. P36930) and imaged using a Nikon confocal microscope.

### Real-time Quantitative Reverse Transcription-Polymerase Chain Reaction for RNA expression

Relative expression of messenger RNA from all *in vitro*, *in vivo* and *ex vivo* experiments were determined by real-time quantitative reverse transcription-polymerase chain reaction (RT-qPCR) on Viia7 1.0 Real-Time PCR system using TaqMan gene expression assays. Reverse transcription with random primers and subsequent PCR were performed with TaqMan RNA-to-CT 1-Step Kit (Applied Biosystems). Raw data for cycle threshold (Ct) values were calculated using the ViiA7 v.1 software with automatically set baseline. The results were analyzed by the ΔΔCt method and GAPDH (Glyceraldehyde 3-phosphate dehydrogenase) was used as a housekeeping gene. Fold change was calculated by taking the average over all the control samples as the baseline. All the probes used in this study were purchased from Thermo Fisher Scientific.

### Connectivity mapping

The connectivity mapping has been used previously by us and others *(22, 32, 72)* and the pipeline utilized here has been recently described*(26)*. Briefly, to obtain compound signatures, A549 and MCF7 cell lines were exposed for 7 hours to two concentrations (3x IC50 and 10x IC50) of compounds made available by AstraZeneca, followed by RNA extraction and RNASeq analysis (single end, read length 49bp at BGI Genomics). A disease library was generated by collating disease signatures from publicly available databases and consisted of 764 signatures representing 310 different diseases. IPF disease signatures were obtained from GSE24206 *(34)* and GSE44723 *(35)* Disease signatures were compared to compound signatures, quantifying the extent of their induced expression change using a modified Kolmogorov-Smirnov score to determine a single connectivity score. The adjusted score reflects the specificity of the connection and allows for better prioritization of disease connections. Connectivity scores were considered significant at FDR < 0.01. Leading edge enrichment analysis was performed on the condition which showed the highest connectivity (FDR 1.1 x 10^-19^), (“Saracatinib_MCF7_Low_Dose” versus “Advanced_IPF_explant_upper_lobe” obtained from GSE24206). Gene sets used in the enrichment analysis were derived from a combination of publicly available sources including Hallmark and KEA *(73)*.

### Disease Enrichment Analysis

For each compound signature, the full list of 764 disease signatures was rank ordered according to ascending connectivity score. For each disease signature in the disease signature library, we collated the relevant Disease Ontology Identifier (DOID), which reflects the disease concept classification that best represents a given signature *(74)*. For each DOID that has at least three corresponding disease signatures, we calculated a signed running sum enrichment score, which reflects whether that DOID is over-represented at the extreme ends of the ranked disease list that has been ordered according to connectivity with a specific compound signature. DOID enrichments with a positive score indicate a disease with multiple individual disease signatures at the top of the ordered list i.e., diseases that are expected to be “normalized” by a given compound. Statistical significance of DEA scores is based on comparison to a distribution of 1000 permuted null scores, generated by calculating scores from randomized DOID sets that contain an equivalent number of disease signatures to the true set being evaluated. Raw p-values are adjusted using the Benjamini-Hochberg method of controlling the false discovery rate.

### RNA sequencing

NHLFs were treated with drugs or vehicle controls with and without TGF-β (10 conditions: 3 replicates each of vehicle A + media; vehicle A + TGF-β; saracatinib + TGF-β; nintedanib + TGF-β; pirfenidone + TGF-β; saracatinib + vehicle B; nintedanib+ vehicle B; pirfenidone + vehicle B; vehicle A + vehicle B; media + vehicle B). Total RNA was extracted with miRNeasy Mini Kit (Qiagen-217004) according to the manufacturer’s instructions. The purity of the RNA was verified using a NanoDrop at 260 nm, and the quality of the RNA was assessed using the Agilent 2100 Bioanalyzer (Agilent Technologies). Libraries were prepared using KAPA Stranded mRNA-Seq Kit. mRNA was enriched by ribosomal RNA depletion. Library quality was checked using Agilent TapeStation analyzer and sequencing was performed with NovaSeq (2X100, 25 M reads per sample). FastQC files were generated and trimmed (https://www.bioinformatics.babraham.ac.uk/projects/trim_galore/) and quality review was performed by FastQC (http://www.bioinformatics.babraham.ac.uk/projects/fastqc/) before and after quality trim. Reads were aligned with STAR aligner (version 2.6.1d, using the default settings) to the HG38 human genome and ENSEMBL94 annotation GTF file. After alignment and summarization with featureCounts of the Subread package (feature Counts release 1.6.3 and picard version 2.18.21), samples were normalized using voom (limma package) and differential expression was carried out with limma *(75)* implemented in R.

Mouse lungs were isolated from bleomycin and TGF-β mice models (8 conditions: 5 replicates each of bleomycin + vehicle, bleomycin + saracatinib, saline + vehicle, saline + saracatinib, TGF-β + vehicle, TGF-β + saracatinib, empty vector + vehicle, empty vector + saracatinib). Total RNA was extracted with miRNeasy Mini Kit (Qiagen-217004) and libraries were prepared as above. Reads were mapped to reference genome mm9 by STAR/3.6.1d *(76)* aligner supplemented with corresponding splice junction file Muc_musculus.GRCm38.96.gtf. At least 90% of reads from each sample were mapped. Duplicates were removed by Picard2.20.2 MarkDuplicates (https://broadinstitute.github.io/picard/), read counts for each gene were obtained by Subread/1.6.3 featureCounts *(77)*, and gene annotations were obtained by biomaRt. Differential expression gene analysis was performed with edgeR *(78)* and limma *(75)* Low-expressed genes were first filtered when counts were no more than 10 in at least 5 samples. After TMM normalization, multidimensional scaling plots were used to explore sample similarities in an unsupervised manner. FDR adjusted p-value < 0.05 was used as the cut-off (Benjamini-Hochberg method).

### Gene set enrichment analysis

Differentially expressed genes in TGF-β treated NHLFs treated with nintedanib and saracatinib were used for pathway enrichment by GSEA (http://software.broadinstitute.org/gsea/msigdb/annotate.jsp). Differentially expressed genes (adj. p-value < 0.05) were included in the enrichment analysis in Fig 2I and Table S2. There were no pathways enriched by nintedanib when including differentially expressed genes with an adjusted p-value < 0.05, therefore, we included genes with an adjusted p-value < 0.1 for both saracatinib and nintedanib for the purposes of this analysis (Figure S4B). The up-regulated and down-regulated gene sets for each drug treatment were used separately. The computed overlaps of each gene set were enriched against the Hallmark and KEGG gene sets from the MSigDB at FDR < 0.05.

Gene set enrichment analysis for both bleomycin and Ad-TGF-β mouse models was carried out using Camera *(79)*, against Hallmark (mouse_H_v5p2.rdata) and KEGG (mouse_c2_v5p2.rdata) datasets. Gene sets with FDR <0.05 were plotted using ggplot2.

### Network analysis of mouse models

Protein-encoding gene interaction network analysis was carried out using Cytoscape v3.7.1 *(80)* with StringAPP plugin *(81)* Breakdown into sub-networks was achieved by clusterMaker plugin MCL cluster function with inflation set at 2.5. Functional enrichment analysis of each cluster was performed using STRING enrichment against a collection of gene set databases including Gene Ontology, KEGG, and Reactome.

### Statistics

For *in vitro* and *in vivo* assays statistical analysis was performed by GraphPad Prism version 8.1.2. Results were analyzed by Mann–Whitney *U* test for comparisons of two groups when sample data were not normally distributed, by unpaired Student’s *t*-test for comparisons of two groups with normal distribution, and by one-way ANOVA with Student–Newman–Keuls *post hoc* test for pairwise comparisons of three or more groups or more than 10 per group. Efficacy experiments were designed for ten-fifteen animals in the control and treated groups, to allow for 82% power to detect a difference of 20% between the two groups at a statistical significance level of 0.05, but the actual size of groups differed because of mortality. All data is presented as (means+ SEM) and the differences were considered statistically significant at *P* < 0.05.

## Acknowledgments

We thank the members of the Yale Center for Genome Analysis who have helped with all aspects of RNA sequencing.

## Author contributions

G.P.D. and N.K and J.D. conceptualized, acquired funding, and supervised the study. L.P.C provided saracatinib and necessary guidance for this project. All *in vitro* studies were performed by F.A., M.N., C.M., K.C., H.M.R., K.W.K., N.B. and Y.A. All *in vivo* studies were performed by F.A., D.F., K.B., K.R., S.D., J.C.S., Q.L., T.B. and N.O. All ex *vivo* studies were performed by F.A. M.C and K.R. RNA sequencing on *in vitro* and *in vivo* models was performed by F.A. and D.F. Transcriptomic data was processed, curated, and visualized by C.B., X.W., B.R. and L.L. The original RNA sequencing to identify the compound signatures was performed by B.M., R.H. and A.B. Connectivity mapping and disease enrichment analysis were performed by C.B. and B.R. The manuscript was drafted by F.A., C.B., G.P.D., N.K. and was reviewed and edited by all other authors.

## Data and materials availability

All data, code, and materials used in the analysis is available to any researcher for purposes of reproducing or extending the analysis. All raw count expression data from *in vitro* and *in vivo* studies were deposited in the Gene Expression Omnibus (GEO) and the accession numbers are GSE 178518 and GSE17845, respectively.

---

**Figure S1.** TGF-β induces Src activation in normal human lung fibroblasts.

**Figure S2.** Evaluation of the effects of two different doses of saracatinib, nintedanib and pirfenidone in TGF-β1-induced profibrotic gene expressions in normal human lung fibroblasts.

**Figure S3.** Saracatinib inhibits TGF-β1–induced phenotypic changes in primary human lung fibroblasts isolated from three different donors to confirm the consistency across donors.

**Figure S4.** Transcriptomic analysis revealed that both nintedanib and saracatinib target several common gene sets.

**Figure S5.** Mice receiving saracatinib treatment recovered their weight losses in both bleomycin and Ad-TGF-β mouse models.

**Figure S6.** Saracatinib inhibits *Col3a1* in bleomycin and Ad-TGF-β mouse models.

**Figure S7.** Representative images (Dorsal view of three-dimensional reconstructions and Axial view) of microCT on mouse lung tissues in bleomycin mice model in the indicated groups.

**Figure S8.** Representative images of lung tissues in Ad-TGF-β mouse model.

**Figure S9.** Western blot analysis of lung tissues identified saracatinib significantly inhibits Phospho-Smad3 signaling in both animal models of pulmonary fibrosis.

**Figure S10.** LIVE/DEAD® Viability/Cytotoxicity identified the viability of PCLS after 5 days in culture.

**Figure S11.** Enlarged version of Cytoscape network analysis of common differentially expressed genes shared by both the bleomycin and Ad-TGF-β mouse models that are reversed by saracatinib.

**Table S1. Connection between IPF and saracatinib signature.**

**Table S2. Enrichment analysis of significantly differentially expressed genes.**

## References

1. L. Richeldi, H. R. Collard, M. G. Jones, Idiopathic pulmonary fibrosis. Lancet 389, 1941–1952 (2017).

2. F. J. Martinez, A. Chisholm, H. R. Collard, K. R. Flaherty, J. Myers, G. Raghu, S. L. Walsh, E. S. White, L. Richeldi, The diagnosis of idiopathic pulmonary fibrosis: current and future approaches. Lancet Respir Med 5, 61–71 (2017).

3. D. J. Lederer, F. J. Martinez, Idiopathic Pulmonary Fibrosis. N Engl J Med 379, 797–798 (2018).

4. G. Raghu, S. Y. Chen, W. S. Yeh, B. Maroni, Q. Li, Y. C. Lee, H. R. Collard, Idiopathic pulmonary fibrosis in US Medicare beneficiaries aged 65 years and older: incidence, prevalence, and survival, 2001-11. Lancet Respir Med 2, 566-572 (2014).

5. A. S. Lee, I. Mira-Avendano, J. H. Ryu, C. E. Daniels, The burden of idiopathic pulmonary fibrosis: an unmet public health need. Respir Med 108, 955–967 (2014).

6. T. M. Maher, E. Bendstrup, L. Dron, J. Langley, G. Smith, J. M. Khalid, H. Patel, M. Kreuter, Global incidence and prevalence of idiopathic pulmonary fibrosis. Respir Res 22, 197 (2021).

7. A. L. Olson, J. J. Swigris, Idiopathic pulmonary fibrosis: diagnosis and epidemiology. Clinics in chest medicine 33, 41–50 (2012).

8. B. M. A. Seibold, A. L. Wise, M. C. Speer, M. P. Steele, K. K. Brown, J. E. Loyd, T. E. Fingerlin, W. Zhang, G. Gudmundsson, S. D. Groshong, C. M. Evans, S. Garantziotis, K.B. Adler, B. F. Dickey, R. M. du Bois, I. V. Yang, A. Herron, D. Kervitsky, J. L. Talbert, C. Markin, J. Park, A. L. Crews, S. H. Slifer, S. Auerbach, M. G. Roy, J. Lin, C. E. Hennessy, M. I. Schwarz, D. A. Schwartz, A common MUC5B promoter polymorphism and pulmonary fibrosis. N Eng J Med 364, 1503–1512 (2011).

9. M. Y. Armanios, J. J. Chen, J. D. Cogan, J. K. Alder, R. G. Ingersoll, C. Markin, W. E. Lawson, M. Xie, I. Vulto, J. A. Phillips, 3rd, P. M. Lansdorp, C. W. Greider, J. E. Loyd, Telomerase mutations in families with idiopathic pulmonary fibrosis. N Engl J Med 356, 1317–1326 (2007).

10. T. E. King, Jr., A. Pardo, M. Selman, Idiopathic pulmonary fibrosis. Lancet 378, 1949–1961 (2011).

11. T. A. Wynn, Integrating mechanisms of pulmonary fibrosis. The Journal of experimental medicine 208, 1339–1350 (2011).

12. V. J. Thannickal, C. A. Henke, J. C. Horowitz, P. W. Noble, J. Roman, P. J. Sime, Y. Zhou, R. G. Wells, E. S. White, D. J. Tschumperlin, Matrix biology of idiopathic pulmonary fibrosis: a workshop report of the national heart, lung, and blood institute. Am J Pathol 184, 1643–1651 (2014).

13. R. Amanchy, J. Zhong, R. Hong, J. H. Kim, M. Gucek, R. N. Cole, H. Molina, A. Pandey, Identification of c-Src tyrosine kinase substrates in platelet-derived growth factor receptor signaling. Mol Oncol 3, 439–450 (2009).

14. G. M. Twamley-Stein, R. Pepperkok, W. Ansorge, S. A. Courtneidge, The Src family tyrosine kinases are required for platelet-derived growth factor-mediated signal transduction in NIH 3T3 cells. Proc Natl Acad Sci U S A 90, 7696–7700 (1993).

15. A. G. Tatosyan, O. A. Mizenina, Kinases of the Src family: structure and functions. Biochemistry (Mosc) 65, 49–58 (2000).

16. M. Hu, P. Che, X. Han, G. Q. Cai, G. Liu, V. Antony, T. Luckhardt, G. P. Siegal, Y. Zhou, R. M. Liu, L. P. Desai, P. J. O’Reilly, V. J. Thannickal, Q. Ding, Therapeutic targeting of SRC kinase in myofibroblast differentiation and pulmonary fibrosis. J Pharmacol Exp Ther 351, 87–95 (2014).

17. G. Hughes, H. Toellner, H. Morris, C. Leonard, N. Chaudhuri, Real World Experiences: Pirfenidone and Nintedanib are Effective and Well Tolerated Treatments for Idiopathic Pulmonary Fibrosis. J Clin Med 5, (2016).

18. L. Richeldi, R. M. du Bois, G. Raghu, A. Azuma, K. K. Brown, U. Costabel, V. Cottin, K. R. Flaherty, D. M. Hansell, Y. Inoue, D. S. Kim, M. Kolb, A. G. Nicholson, P. W. Noble, M. Selman, H. Taniguchi, M. Brun, F. Le Maulf, M. Girard, S. Stowasser, R. Schlenker-Herceg, B. Disse, H. R. Collard, I. T. Investigators, Efficacy and safety of nintedanib in idiopathic pulmonary fibrosis. N Engl J Med 370, 2071–2082 (2014).

19. T. E. King, Jr., W. Z. Bradford, S. Castro-Bernardini, E. A. Fagan, I. Glaspole, M. K. Glassberg, E. Gorina, P. M. Hopkins, D. Kardatzke, L. Lancaster, D. J. Lederer, S. D. Nathan, C. A. Pereira, S. A. Sahn, R. Sussman, J. J. Swigris, P. W. Noble, A. S. Group, A phase 3 trial of pirfenidone in patients with idiopathic pulmonary fibrosis. N Engl J Med 370, 2083–2092 (2014).

20. T. Corte, F. Bonella, B. Crestani, M. G. Demedts, L. Richeldi, C. Coeck, K. Pelling, M. Quaresma, J. A. Lasky, Safety, tolerability and appropriate use of nintedanib in idiopathic pulmonary fibrosis. Respir Res 16, 116 (2015).

21. F. Varone, G. Sgalla, B. Iovene, T. Bruni, L. Richeldi, Nintedanib for the treatment of idiopathic pulmonary fibrosis. Expert Opin Pharmacother 19, 167–175 (2018).

22. J. T. Dudley, M. Sirota, M. Shenoy, R. K. Pai, S. Roedder, A. P. Chiang, A. A. Morgan, M. M. Sarwal, P. J. Pasricha, A. J. Butte, Computational repositioning of the anticonvulsant topiramate for inflammatory bowel disease. Sci Transl Med 3, 96ra76 (2011).

23. M. Sirota, J. T. Dudley, J. Kim, A. P. Chiang, A. A. Morgan, A. Sweet-Cordero, J. Sage, A. J. Butte, Discovery and preclinical validation of drug indications using compendia of public gene expression data. Sci Transl Med 3, 96ra77 (2011).

24. A. M. Brum, J. van de Peppel, C. S. van der Leije, M. Schreuders-Koedam, M. Eijken, B. C. van der Eerden, J. P. van Leeuwen, Connectivity Map-based discovery of parbendazole reveals targetable human osteogenic pathway. Proc Natl Acad Sci U S A 112, 12711–12716 (2015).

25. V. van Noort, S. Scholch, M. Iskar, G. Zeller, K. Ostertag, C. Schweitzer, K. Werner, J. Weitz, M. Koch, P. Bork, Novel drug candidates for the treatment of metastatic colorectal cancer through global inverse gene-expression profiling. Cancer Res 74, 5690–5699 (2014).

26. D. Bhattacharya, C. Becker, B. Readhead, N. Goossens, J. Novik, M. I. Fiel, L. P. Cousens, B. Magnusson, A. Backmark, R. Hicks, J. T. Dudley, S. L. Friedman, Repositioning of a novel GABA-B receptor agonist, AZD3355 (Lesogaberan), for the treatment of non-alcoholic steatohepatitis. Sci Rep 11, 20827 (2021).

27. Y. M. Chang, L. Bai, S. Liu, J. C. Yang, H. J. Kung, C. P. Evans, Src family kinase oncogenic potential and pathways in prostate cancer as revealed by AZD0530. Oncogene 27, 6365–6375 (2008).

28. T. P. Green, M. Fennell, R. Whittaker, J. Curwen, V. Jacobs, J. Allen, A. Logie, J. Hargreaves, D. M. Hickinson, R. W. Wilkinson, P. Elvin, B. Boyer, N. Carragher, P. A. Ple, A. Bermingham, G. A. Holdgate, W. H. Ward, L. F. Hennequin, B. R. Davies, G. F. Costello, Preclinical anticancer activity of the potent, oral Src inhibitor AZD0530. Mol Oncol 3, 248–261 (2009).

29. J. Baselga, A. Cervantes, E. Martinelli, I. Chirivella, K. Hoekman, H. I. Hurwitz, D. I. Jodrell, P. Hamberg, E. Casado, P. Elvin, A. Swaisland, R. Iacona, J. Tabernero, Phase I safety, pharmacokinetics, and inhibition of SRC activity study of saracatinib in patients with solid tumors. Clin Cancer Res 16, 4876–4883 (2010).

30. F. Ahangari, C. Becker, D. Foster, M. Chioccioli, M. Nelson, K. Beke, C. Meador, X. Wang, K. Corell, H. Roybal, G. Deluliis, K. Rose, J. C. Schupp, Q. Li, T. Adams, Y. Aschner, L. Lilli, B. Readhead, L. Cousens, J. Dudley, N. Kaminski, G. P. Downey, Saracatinib Is a Potential Novel Therapeutic for Pulmonary Fibrosis. Am J Resp Crit Care 201, (2020).

31. J. Lamb, E. D. Crawford, D. Peck, J. W. Modell, I. C. Blat, M. J. Wrobel, J. Lerner, J. P. Brunet, A. Subramanian, K. N. Ross, M. Reich, H. Hieronymus, G. Wei, S. A. Armstrong, S. J. Haggarty, P. A. Clemons, R. Wei, S. A. Carr, E. S. Lander, T. R. Golub, The Connectivity Map: using gene-expression signatures to connect small molecules, genes, and disease. Science 313, 1929–1935 (2006).

32. A. Subramanian, R. Narayan, S. M. Corsello, D. D. Peck, T. E. Natoli, X. Lu, J. Gould, J. F. Davis, A. A. Tubelli, J. K. Asiedu, D. L. Lahr, J. E. Hirschman, Z. Liu, M. Donahue, B. Julian, M. Khan, D. Wadden, I. C. Smith, D. Lam, A. Liberzon, C. Toder, M. Bagul, M. Orzechowski, O. M. Enache, F. Piccioni, S. A. Johnson, N. J. Lyons, A. H. Berger, A. F. Shamji, A. N. Brooks, A. Vrcic, C. Flynn, J. Rosains, D. Y. Takeda, R. Hu, D. Davison, J. Lamb, K. Ardlie, L. Hogstrom, P. Greenside, N. S. Gray, P. A. Clemons, S. Silver, X. Wu, W. N. Zhao, W. Read-Button, X. Wu, S. J. Haggarty, L. V. Ronco, J. S. Boehm, S. L. Schreiber, J. G. Doench, J. A. Bittker, D. E. Root, B. Wong, T. R. Golub, A Next Generation Connectivity Map: L1000 Platform and the First 1,000,000 Profiles. Cell 171, 1437–1452 e1417 (2017).

33. Z. Wang, C. D. Monteiro, K. M. Jagodnik, N. F. Fernandez, G. W. Gundersen, A. D. Rouillard, S. L. Jenkins, A. S. Feldmann, K. S. Hu, M. G. McDermott, Q. Duan, N. R. Clark, M. R. Jones, Y. Kou, T. Goff, H. Woodland, F. M. R. Amaral, G. L. Szeto, O. Fuchs, S. M. Schussler-Fiorenza Rose, S. Sharma, U. Schwartz, X. B. Bausela, M. Szymkiewicz, V. Maroulis, A. Salykin, C. M. Barra, C. D. Kruth, N. J. Bongio, V. Mathur, R. D. Todoric, U. E. Rubin, A. Malatras, C. T. Fulp, J. A. Galindo, R. Motiejunaite, C. Juschke, P. C. Dishuck, K. Lahl, M. Jafari, S. Aibar, A. Zaravinos, L. H. Steenhuizen, L. R. Allison, P. Gamallo, F. de Andres Segura, T. Dae Devlin, V. Perez-Garcia, A. Ma’ayan, Extraction and analysis of signatures from the Gene Expression Omnibus by the crowd. Nat Commun 7, 12846 (2016).

34. E. B. Meltzer, W. T. Barry, T. A. D’Amico, R. D. Davis, S. S. Lin, M. W. Onaitis, L. D. Morrison, T. A. Sporn, M. P. Steele, P. W. Noble, Bayesian probit regression model for the diagnosis of pulmonary fibrosis: proof-of-principle. BMC Med Genomics 4, 70 (2011).

35. R. Peng, S. Sridhar, G. Tyagi, J. E. Phillips, R. Garrido, P. Harris, L. Burns, L. Renteria, J. Woods, L. Chen, J. Allard, P. Ravindran, H. Bitter, Z. Liang, C. M. Hogaboam, C. Kitson, D. C. Budd, J. S. Fine, C. M. Bauer, C. S. Stevenson, Bleomycin induces molecular changes directly relevant to idiopathic pulmonary fibrosis: a model for “active” disease. PLoS One 8, e59348 (2013).

36. J. Xiong, J. S. Wu, S. S. Mao, X. N. Yu, X. X. Huang, Effect of saracatinib on pulmonary metastases from hepatocellular carcinoma. Oncol Rep 36, 1483–1490 (2016).

37. J. Jin, S. Togo, K. Kadoya, M. Tulafu, Y. Namba, M. Iwai, J. Watanabe, K. Nagahama, T. Okabe, M. Hidayat, Y. Kodama, H. Kitamura, T. Ogura, N. Kitamura, K. Ikeo, S. Sasaki, S. Tominaga, K. Takahashi, Pirfenidone attenuates lung fibrotic fibroblast responses to transforming growth factor-beta1. Respir Res 20, 119 (2019).

38. J. Huang, C. Beyer, K. Palumbo-Zerr, Y. Zhang, A. Ramming, A. Distler, K. Gelse, O. Distler, G. Schett, L. Wollin, J. H. Distler, Nintedanib inhibits fibroblast activation and ameliorates fibrosis in preclinical models of systemic sclerosis. Ann Rheum Dis 75, 883–890 (2016).

39. L. Wollin, D. Togbe, B. Ryffel, Effects of Nintedanib in an Animal Model of Liver Fibrosis. Biomed Res Int 2020, 3867198 (2020).

40. S. I. Rothschild, Clinical potential of nintedanib for the second-line treatment of advanced non-small-cell lung cancer: current evidence. Lung Cancer (Auckl) 5, 51–57 (2014).

41. K. E. Hostettler, J. Zhong, E. Papakonstantinou, G. Karakiulakis, M. Tamm, P. Seidel, Q. Sun, J. Mandal, D. Lardinois, C. Lambers, M. Roth, Anti-fibrotic effects of nintedanib in lung fibroblasts derived from patients with idiopathic pulmonary fibrosis. Respir Res 15, 157 (2014).

42. C. J. Schaefer, D. W. Ruhrmund, L. Pan, S. D. Seiwert, K. Kossen, Antifibrotic activities of pirfenidone in animal models. Eur Respir Rev 20, 85–97 (2011).

43. Y. Y. Lu, X. K. Zhao, L. Yu, F. Qi, B. Zhai, C. Q. Gao, Q. Ding, Interaction of Src and Alpha-V Integrin Regulates Fibroblast Migration and Modulates Lung Fibrosis in A Preclinical Model of Lung Fibrosis. Sci Rep 7, 46357 (2017).

44. B. B. Moore, C. M. Hogaboam, Murine models of pulmonary fibrosis. Am J Physiol Lung Cell Mol Physiol 294, L152–160 (2008).

45. P. J. Sime, Z. Xing, F. L. Graham, K. G. Csaky, J. Gauldie, Adenovector-mediated gene transfer of active transforming growth factor-beta1 induces prolonged severe fibrosis in rat lung. J Clin Invest 100, 768–776 (1997).

46. N. Frangogiannis, Transforming growth factor-beta in tissue fibrosis. The Journal of experimental medicine 217, e20190103 (2020).

47. K. C. Flanders, Smad3 as a mediator of the fibrotic response. Int J Exp Pathol 85, 47–64 (2004).

48. M. Lehmann, L. Buhl, H. N. Alsafadi, S. Klee, S. Hermann, K. Mutze, C. Ota, M. Lindner, J. Behr, A. Hilgendorff, D. E. Wagner, M. Konigshoff, Differential effects of Nintedanib and Pirfenidone on lung alveolar epithelial cell function in ex vivo murine and human lung tissue cultures of pulmonary fibrosis. Respir Res 19, 175 (2018).

49. M. Cedilak, M. Banjanac, D. Belamaric, A. Paravic Radicevic, I. Faraho, K. Ilic, S. Cuzic, I. Glojnaric, V. Erakovic Haber, M. Bosnar, Precision-cut lung slices from bleomycin treated animals as a model for testing potential therapies for idiopathic pulmonary fibrosis. Pulm Pharmacol Ther 55, 75–83 (2019).

50. G. Yu, G. H. Ibarra, N. Kaminski, Fibrosis: Lessons from OMICS analyses of the human lung. Matrix Biol 68-69, 422-434 (2018).

51. M. G. Palfreyman, Human tissue in target identification and drug discovery. Drug Discov Today 7, 407–409 (2002).

52. N. Shanks, R. Greek, J. Greek, Are animal models predictive for humans? Philos Ethics Humanit Med 4, 2 (2009).

53. A. Musa, L. S. Ghoraie, S. D. Zhang, G. Glazko, O. Yli-Harja, M. Dehmer, B. Haibe-Kains, F. Emmert-Streib, A review of connectivity map and computational approaches in pharmacogenomics. Brief Bioinform 18, 903 (2017).

54. H. Li, C. Zhao, Y. Tian, J. Lu, G. Zhang, S. Liang, D. Chen, X. Liu, W. Kuang, M. Zhu, Src family kinases and pulmonary fibrosis: A review. Biomed Pharmacother 127, 110183 (2020).

55. A. Ulsamer, Y. Wei, K. K. Kim, K. Tan, S. Wheeler, Y. Xi, R. S. Thies, H. A. Chapman, Axin pathway activity regulates in vivo pY654-beta-catenin accumulation and pulmonary fibrosis. J Biol Chem 287, 5164–5172 (2012).

56. Y. Aschner, A. P. Khalifah, N. Briones, C. Yamashita, L. Dolgonos, S. K. Young, M. N. Campbell, D. W. Riches, E. F. Redente, W. J. Janssen, P. M. Henson, J. Sap, N. Vacaresse, A. Kapus, C. A. McCulloch, R. L. Zemans, G. P. Downey, Protein tyrosine phosphatase alpha mediates profibrotic signaling in lung fibroblasts through TGF-beta responsiveness. Am J Pathol 184, 1489–1502 (2014).

57. Y. Aschner, M. Nelson, M. Brenner, H. Roybal, K. Beke, C. Meador, D. Foster, K. A. Correll, P. R. Reynolds, K. Anderson, E. F. Redente, J. Matsuda, D. W. H. Riches, S. D. Groshong, A. Pozzi, J. Sap, Q. Wang, D. Rajshankar, C. A. G. McCulloch, R. L. Zemans, G. P. Downey, Protein tyrosine phosphatase-alpha amplifies transforming growth factor-beta-dependent profibrotic signaling in lung fibroblasts. Am J Physiol Lung Cell Mol Physiol 319, L294–L311 (2020).

58. T. J. Boggon, M. J. Eck, Structure and regulation of Src family kinases. Oncogene 23, 7918–7927 (2004).

59. A. Tzouvelekis, G. Yu, C. L. Lino Cardenas, J. D. Herazo-Maya, R. Wang, T. Woolard, Y. Zhang, K. Sakamoto, H. Lee, J. S. Yi, G. DeIuliis, N. Xylourgidis, F. Ahangari, P. J. Lee, V. Aidinis, E. L. Herzog, R. Homer, A. M. Bennett, N. Kaminski, SH2 Domain-Containing Phosphatase-2 Is a Novel Antifibrotic Regulator in Pulmonary Fibrosis. Am J Respir Crit Care Med 195, 500–514 (2017).

60. Y. Aschner, G. P. Downey, Transforming Growth Factor-beta: Master Regulator of the Respiratory System in Health and Disease. Am J Respir Cell Mol Biol 54, 647–655 (2016).

61. N. Kaminski, J. D. Allard, J. F. Pittet, F. Zuo, M. J. Griffiths, D. Morris, X. Huang, D. Sheppard, R. A. Heller, Global analysis of gene expression in pulmonary fibrosis reveals distinct programs regulating lung inflammation and fibrosis. Proc Natl Acad Sci U S A 97, 1778–1783 (2000).

62. A. Abdollahi, M. Li, G. Ping, C. Plathow, S. Domhan, F. Kiessling, L. B. Lee, G. McMahon, H. J. Grone, K. E. Lipson, P. E. Huber, Inhibition of platelet-derived growth factor signaling attenuates pulmonary fibrosis. The Journal of experimental medicine 201, 925–935 (2005).

63. L. Veracini, M. Franco, A. Boureux, V. Simon, S. Roche, C. Benistant, Two functionally distinct pools of Src kinases for PDGF receptor signalling. Biochem Soc Trans 33, 1313–1315 (2005).

64. H. Xiao, X. Huang, S. Wang, Z. Liu, R. Dong, D. Song, H. Dai, Metformin ameliorates bleomycin-induced pulmonary fibrosis in mice by suppressing IGF-1. Am J Transl Res 12, 940–949 (2020).

65. R. D. Guzy, L. Li, C. Smith, S. J. Dorry, H. Y. Koo, L. Chen, D. M. Ornitz, Pulmonary fibrosis requires cell-autonomous mesenchymal fibroblast growth factor (FGF) signaling. J Biol Chem 292, 10364–10378 (2017).

66. M. Imamura, J. S. Moon, K. P. Chung, K. Nakahira, T. Muthukumar, R. Shingarev, S. W. Ryter, A. M. Choi, M. E. Choi, RIPK3 promotes kidney fibrosis via AKT-dependent ATP citrate lyase. JCI Insight 3, (2018).

67. D. J. Webb, K. Donais, L. A. Whitmore, S. M. Thomas, C. E. Turner, J. T. Parsons, A. F. Horwitz, FAK-Src signalling through paxillin, ERK and MLCK regulates adhesion disassembly. Nat Cell Biol 6, 154–161 (2004).

68. D. V. Pechkovsky, A. K. Scaffidi, T. L. Hackett, J. Ballard, F. Shaheen, P. J. Thompson, V. J. Thannickal, D. A. Knight, Transforming growth factor beta1 induces alphavbeta3 integrin expression in human lung fibroblasts via a beta3 integrin-, c-Src-, and p38 MAPK-dependent pathway. J Biol Chem 283, 12898–12908 (2008).

69. H. N. Alsafadi, C. A. Staab-Weijnitz, M. Lehmann, M. Lindner, B. Peschel, M. Konigshoff, D. E. Wagner, An ex vivo model to induce early fibrosis-like changes in human precision-cut lung slices. Am J Physiol Lung Cell Mol Physiol 312, L896–L902 (2017).

70. F. E. Uhl, S. Vierkotten, D. E. Wagner, G. Burgstaller, R. Costa, I. Koch, M. Lindner, S. Meiners, O. Eickelberg, M. Konigshoff, Preclinical validation and imaging of Wnt-induced repair in human 3D lung tissue cultures. Eur Respir J 46, 1150–1166 (2015).

71. Y. Liu, A. Keikhosravi, G. S. Mehta, C. R. Drifka, K. W. Eliceiri, Methods for Quantifying Fibrillar Collagen Alignment. Methods Mol Biol 1627, 429–451 (2017).

72. J. T. Dudley, R. Tibshirani, T. Deshpande, A. J. Butte, Disease signatures are robust across tissues and experiments. Mol Syst Biol 5, 307 (2009).

73. A. Lachmann, A. Ma’ayan, KEA: kinase enrichment analysis. Bioinformatics 25, 684–686 (2009).

74. L. M. Schriml, E. Mitraka, J. Munro, B. Tauber, M. Schor, L. Nickle, V. Felix, L. Jeng, C. Bearer, R. Lichenstein, K. Bisordi, N. Campion, B. Hyman, D. Kurland, C. P. Oates, S. Kibbey, P. Sreekumar, C. Le, M. Giglio, C. Greene, Human Disease Ontology 2018 update: classification, content and workflow expansion. Nucleic Acids Res 47, D955–D962 (2019).

75. M. E. Ritchie, B. Phipson, D. Wu, Y. Hu, C. W. Law, W. Shi, G. K. Smyth, limma powers differential expression analyses for RNA-sequencing and microarray studies. Nucleic Acids Res 43, e47 (2015).

76. A. Dobin, C. A. Davis, F. Schlesinger, J. Drenkow, C. Zaleski, S. Jha, P. Batut, M. Chaisson, T. R. Gingeras, STAR: ultrafast universal RNA-seq aligner. Bioinformatics 29, 15–21 (2013).

77. Y. Liao, G. K. Smyth, W. Shi, featureCounts: an efficient general purpose program for assigning sequence reads to genomic features. Bioinformatics 30, 923–930 (2014).

78. M. D. Robinson, D. J. McCarthy, G. K. Smyth, edgeR: a Bioconductor package for differential expression analysis of digital gene expression data. Bioinformatics 26, 139–140 (2010).

79. D. Wu, G. K. Smyth, Camera: a competitive gene set test accounting for inter-gene correlation. Nucleic Acids Res 40, e133 (2012).

80. P. Shannon, A. Markiel, O. Ozier, N. S. Baliga, J. T. Wang, D. Ramage, N. Amin, B. Schwikowski, T. Ideker, Cytoscape: a software environment for integrated models of biomolecular interaction networks. Genome Res 13, 2498–2504 (2003).

81. N. T. Doncheva, J. H. Morris, J. Gorodkin, L. J. Jensen, Cytoscape StringApp: Network Analysis and Visualization of Proteomics Data. J Proteome Res 18, 623–632 (2019).

