## Supplementary Materials for "Saracatinib, a Selective Src Kinase Inhibitor, Blocks Fibrotic Responses in *In Vitro, In Vivo* and *Ex Vivo* Models of Pulmonary Fibrosis"

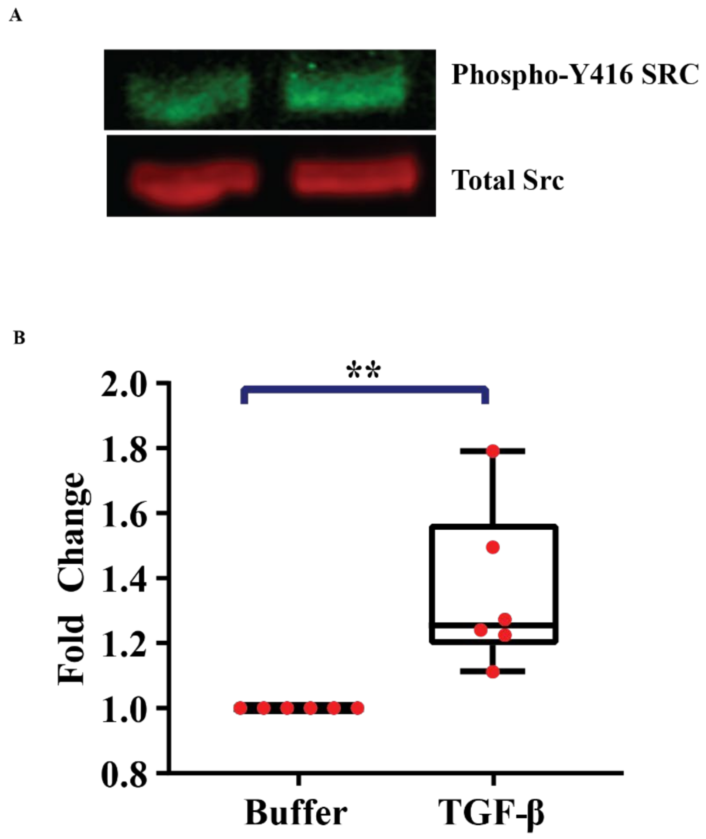


**Figure S1. TGF-β induces Src activation in normal human lung fibroblasts.**

Primary human lung fibroblasts were plated on rat tail collagen-coated non-tissue culture plastic, serum starved overnight, and then stimulated with human recombinant TGF-β (2 ng/ml) for 60 min at 37°C. Protein was solubilized in RIPA buffer and then subject to SDS-PAGE and immunoblotting with antibodies to total Src and phospo-Y416 Src. (A) Representative image of the western blot. (B) Densitometry of the western blots. (n=6), ** P=0.0055.

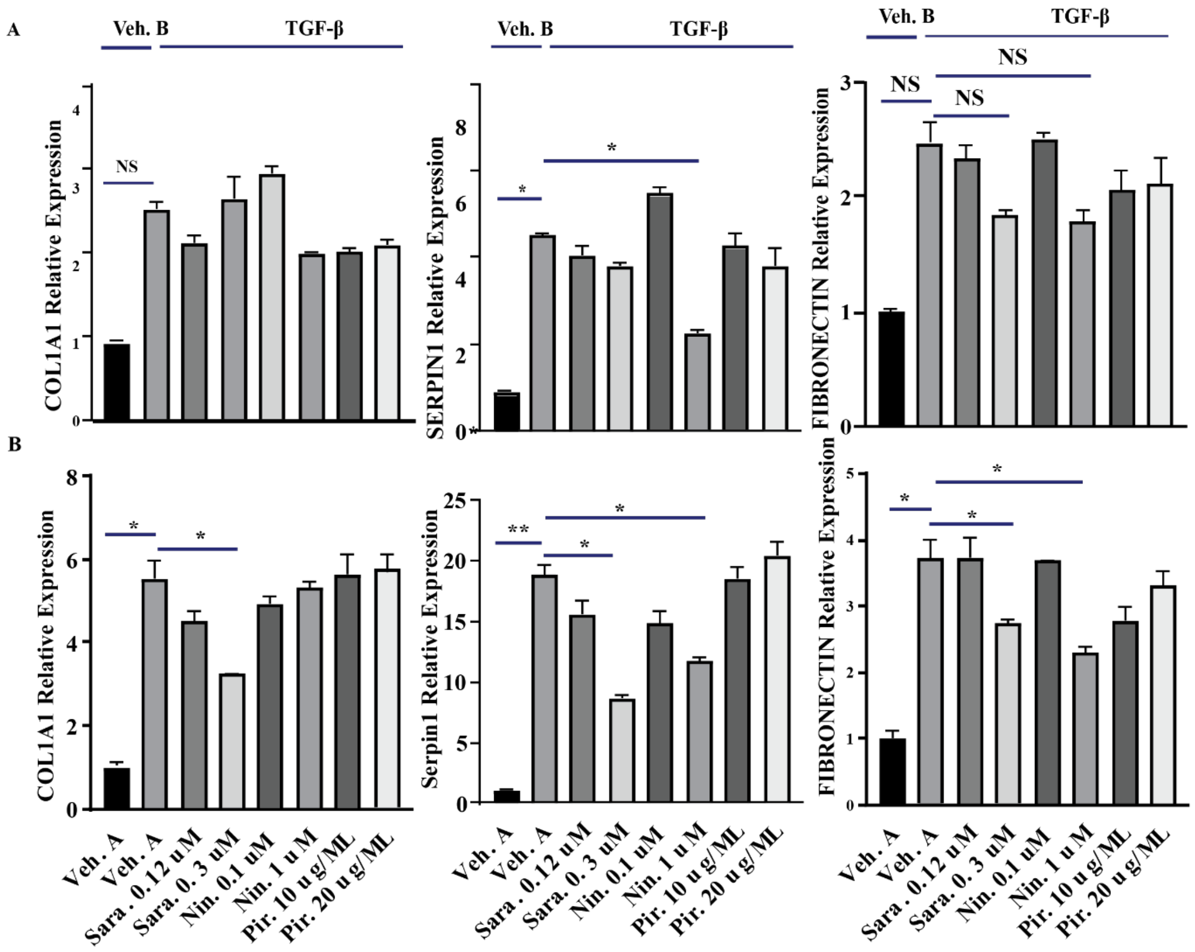


**Figure S2. Evaluation of the effects of two different doses of saracatinib, nintedanib and pirfenidone in TGF-β1-induced profibrotic gene expressions in normal human lung fibroblasts.**

Cells were serum-starved overnight and then incubated with inhibitors at two selected clinically relevant doses (saracatinib: (0.12 & 0.3 μM), nintedanib: (0.1 &1 μM), pirfenidone: (10 & 20 μg/ml) or vehicle A (DMSO)) for 60 min followed by stimulation with human recombinant TGF-β1 (2 ng/ml) or vehicle B as control for 6 and 24 hours. (A) 6-hour timepoint; RT-qPCR analysis for *COL1A1*, *SERPIN-1* (*Pie-1*) and *FIBRONECTIN* in the indicated treatment groups. (B) 24-

hour timepoint; RT-qPCR analysis for *COL1A1*, *SERPIN-1* (*Pie-1*) and *FIBRONECTIN* in the indicated treatment groups. All data is presented as means+ SEM, *P<0.05, **P<0.01, (n=4).

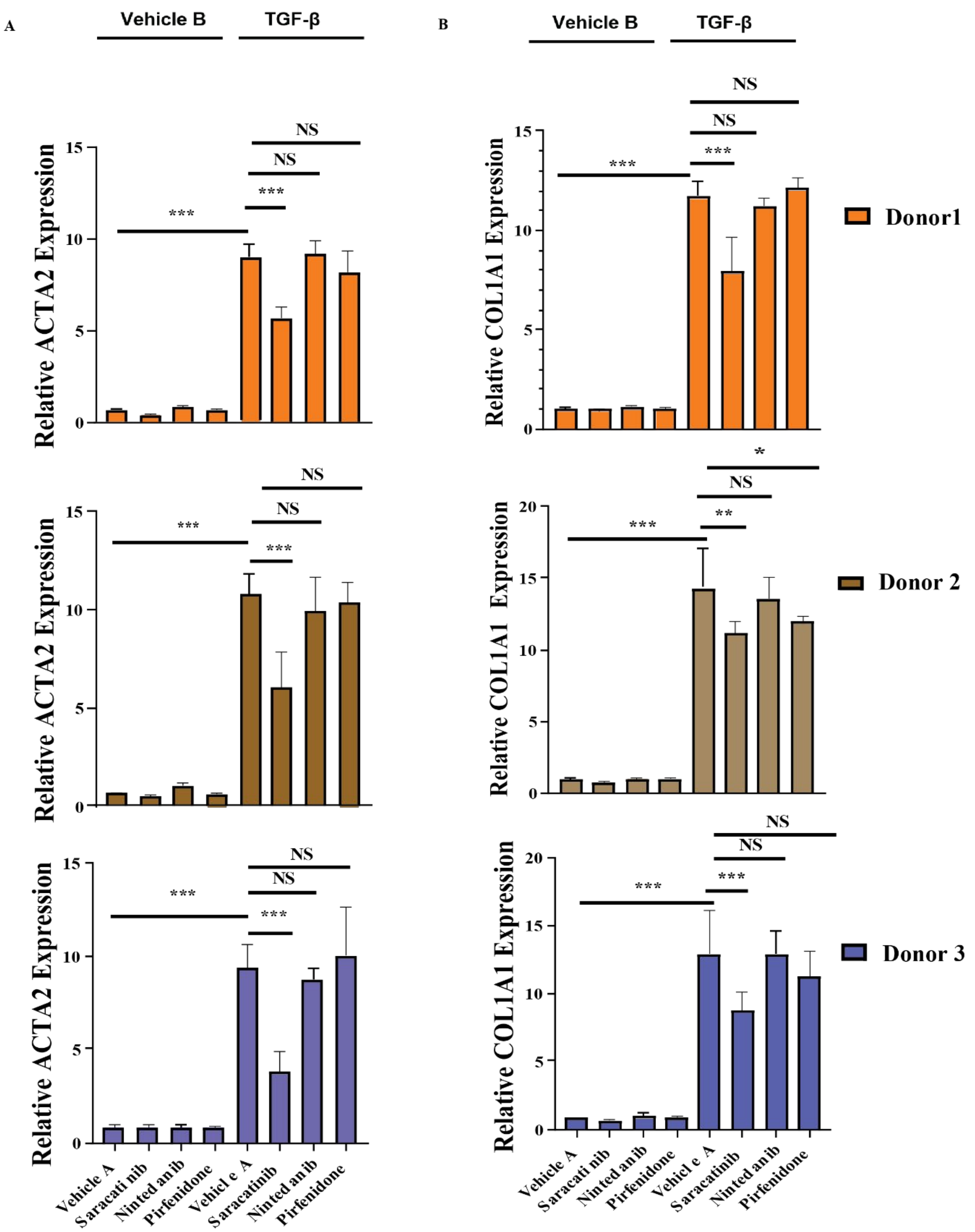
A
Vehicle B
TGF-β
B
Vehicle B
TGF-β
NS
NS
***
***
Relative ACTA2 Expression
Relative COL1A1 Expression
Donor1
Donor 2
Donor 3
*
**
Vehicle A
Saracati nib
Ninted anib
Pirfenidone
Vehicl e A
Saracatinib
Ninted anib
Pirfenidone

**Figure S3. Saracatinib inhibits TGF-β1–induced phenotypic changes in primary human lung fibroblasts isolated from three different donors to confirm the consistency across donors.** (A and B) RT-qPCR analysis for (A) *ACTA2* (B) *COL1A1* on primary human lung fibroblasts isolated from three different donors in the indicated treatment groups. All experiments were performed independently. (means+ SEM), *P<0.05, **P<0.01, ***P<0.001, (n=6).

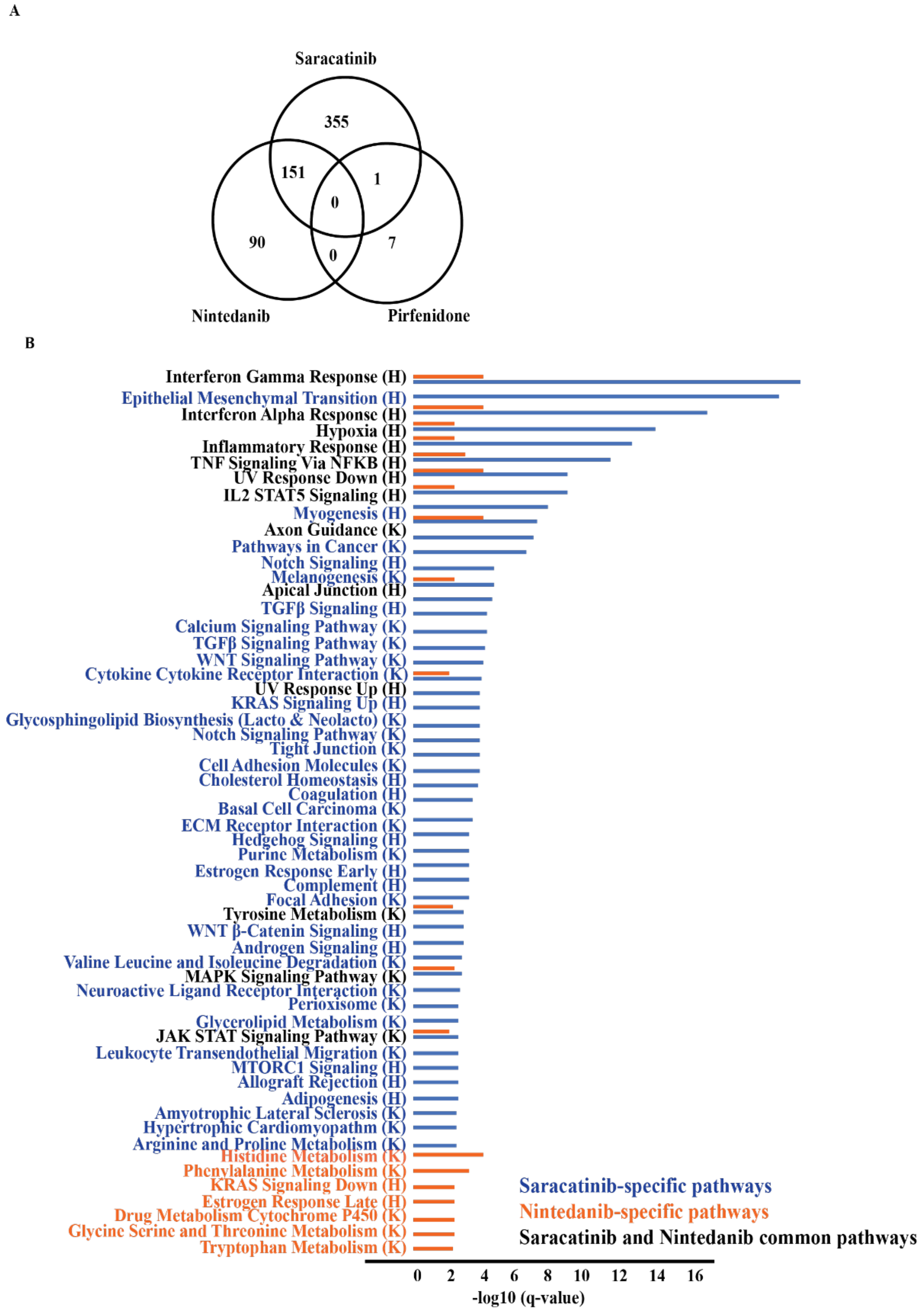
A
Saracatinib
355
151
1
0
90
0
7
Nintedanib
Pirfenidone
B
Interferon Gamma Response (H)
Epithelial Mesenchymal Transition (H)
Interferon Alpha Response (H)
Hypoxia (H)
Inflammatory Response (H)
TNF Signaling Via NFKB (H)
UV Response Down (H)
IL2 STAT5 Signaling (H)
Myogenesis (H)
Axon Guidance (K)
Pathways in Cancer (K)
Notch Signaling (H)
Melanogenesis (K)
Apical Junction (H)
TGFβ Signaling (H)
Calcium Signaling Pathway (K)
TGFβ Signaling Pathway (K)
WNT Signaling Pathway (K)
Cytokine Cytokine Receptor Interaction (K)
UV Response Up (H)
KRAS Signaling Up (H)
Glycosphingolipid Biosynthesis (Lacto & Neolacto) (K)
Notch Signaling Pathway (K)
Tight Junction (K)
Cell Adhesion Molecules (K)
Cholesterol Homeostasis (H)
Coagulation (H)
Basal Cell Carcinoma (K)
ECM Receptor Interaction (K)
Hedgehog Signaling (H)
Purine Metabolism (K)
Estrogen Response Early (H)
Complement (H)
Focal Adhesion (K)
Tyrosine Metabolism (K)
WNT β-Catenin Signaling (H)
Androgen Signaling (H)
Valine Leucine and Isoleucine Degradation (K)
MAPK Signaling Pathway (K)
Neuroactive Ligand Receptor Interaction (K)
Perioxisome (K)
Glycerolipid Metabolism (K)
JAK STAT Signaling Pathway (K)
Leukocyte Transendothelial Migration (K)
MTORC1 Signaling (H)
Allograft Rejection (H)
Adipogenesis (H)
Amyotrophic Lateral Sclerosis (K)
Hypertrophic Cardiomyopathm (K)
Arginine and Proline Metabolism (K)
Histidine Metabolism (K)
Phenylalanine Metabolism (K)
KRAS Signaling Down (H)
Estrogen Response Late (H)
Drug Metabolism Cytochrome P450 (K)
Glycine Serine and Threonine Metabolism (K)
Tryptophan Metabolism (K)
Saracatinib-specific pathways
Nintedanib-specific pathways
Saracatinib and Nintedanib common pathways
0 2 4 6 8 10 12 14 16
-log10 (q-value)

**Figure S4. Transcriptomic analysis revealed that both nintedanib and saracatinib target several common gene sets**.

(A) Comparison of number of genes that are differentially expressed following treatment by saracatinib, nintedanib or pirfenidone in TGF-treated cells (FDR <0.05). Of the 507 genes that were significantly differentially expressed in the presence of saracatinib, 355 were unique to saracatinib. Of the 241 genes that were differentially expressed in the presence of nintedanib, 90 were unique to nintedanib. Eight genes were differentially expressed in the presence of pirfenidone, and of these seven were unique. Saracatinib and pirfenidone share one gene in common (ZNF214); saracatinib and nintedanib share 151 genes in common. (B) Functional enrichment of significantly differentially expressed genes (FDR < 0.1, *no enrichments were identified for nintedanib when genes with FDR <0.05 were used) in response to saracatinib (top 50 gene sets shown; blue) or nintedanib (all gene sets; orange). Common gene sets enriched by both saracatinib and nintedanib are shown in black. All gene sets shown are significant at FDR <0.05 and are from Hallmark (H) or Kegg (K).

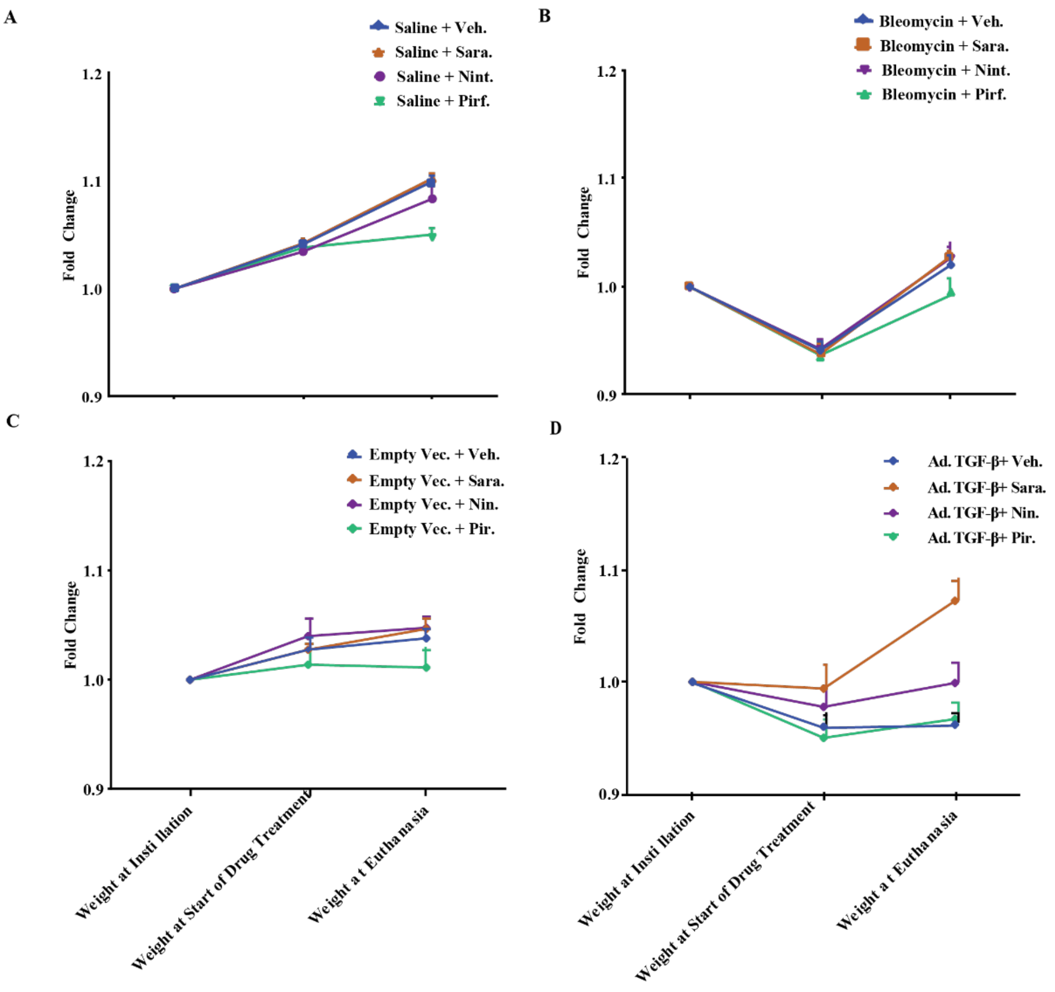


**Figure S5. Mice receiving saracatinib treatment recovered their weight losses in both bleomycin and Ad-TGF-β mouse models.**

(A) Saline treated mice (B) bleomycin treated mice (C) Empty vector treated mice (D) Ad-TGF-β treated mice. All mice were weighed daily. All data is shown as fold change in body weight, (n=6 in saline and empty vector treated and n≥12 in Bleomycin and Ad-TGF-β treated groups).

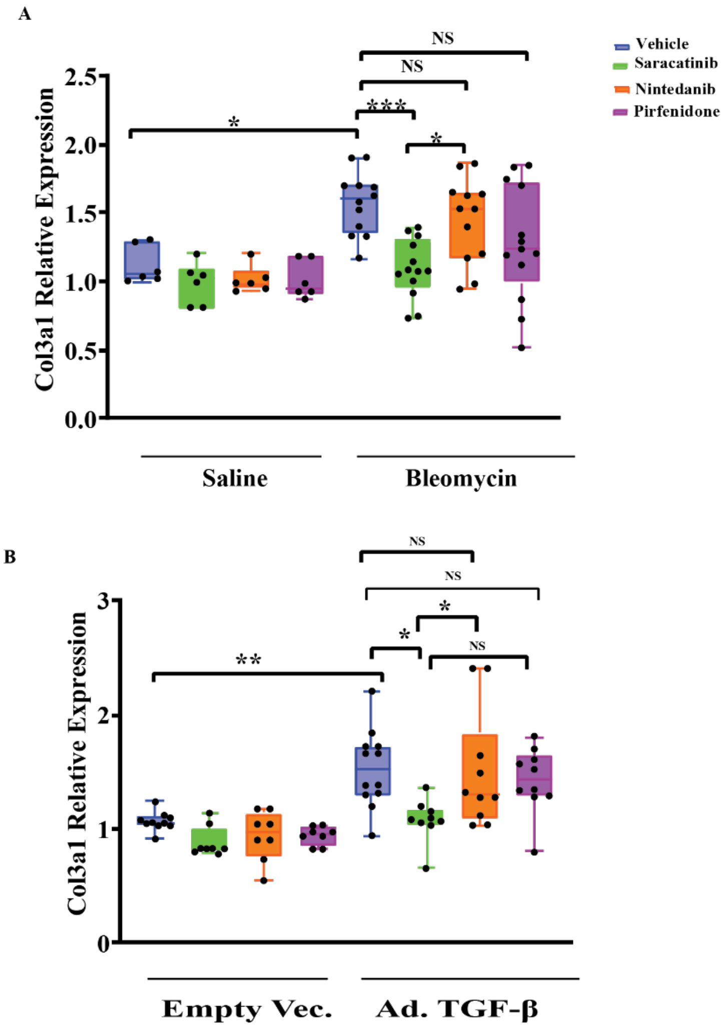


**Figure S6**. **Saracatinib inhibits *Col3a1* in bleomycin and Ad-TGF-β mouse models.**

RT-qPCR analysis of mice lungs in the indicated treatment groups, (A) in bleomycin-induced lung fibrosis model (B) in Ad-TGF-β -induced lung fibrosis model. All data is presented as means+ SEM, *P<0.05, **P<0.01, ***P<0.001, (n=6 in saline and n≥12 in bleomycin and Ad-TGF-β treated groups).

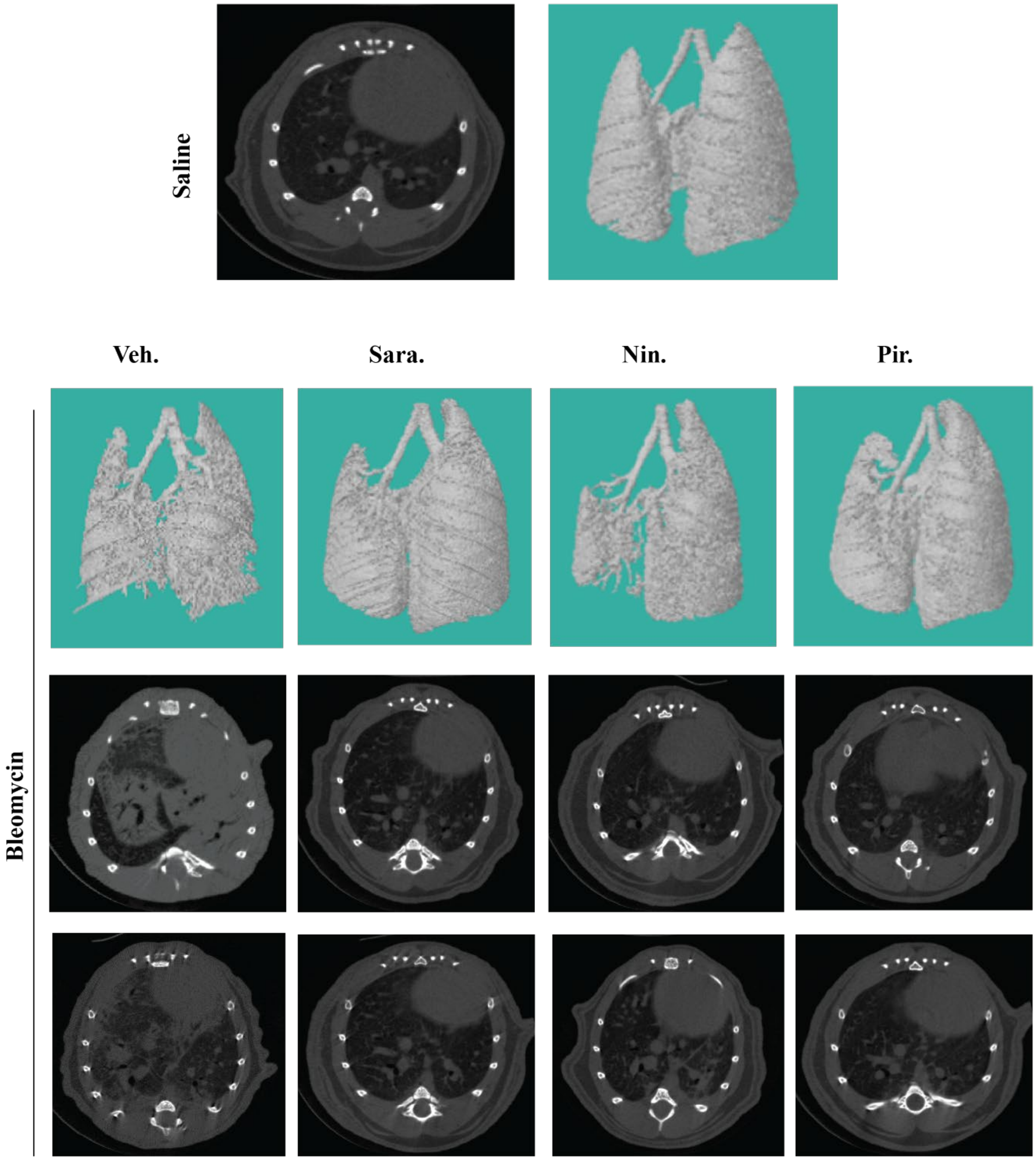
Saline
Veh.
Sara.
Nin.
Pir.
Bleomycin

**Figure S7. Representative images (Dorsal view of three-dimensional reconstructions and Axial view) of microCT on mouse lung tissues in bleomycin mice model in the indicated groups.**

Gross abnormality resulting from bleomycin- induced lung fibrosis is alleviated after treatment, while saracatinib and nintedanib significantly attenuated the bleomycin-induced radiographic alterations in lung parenchyma.

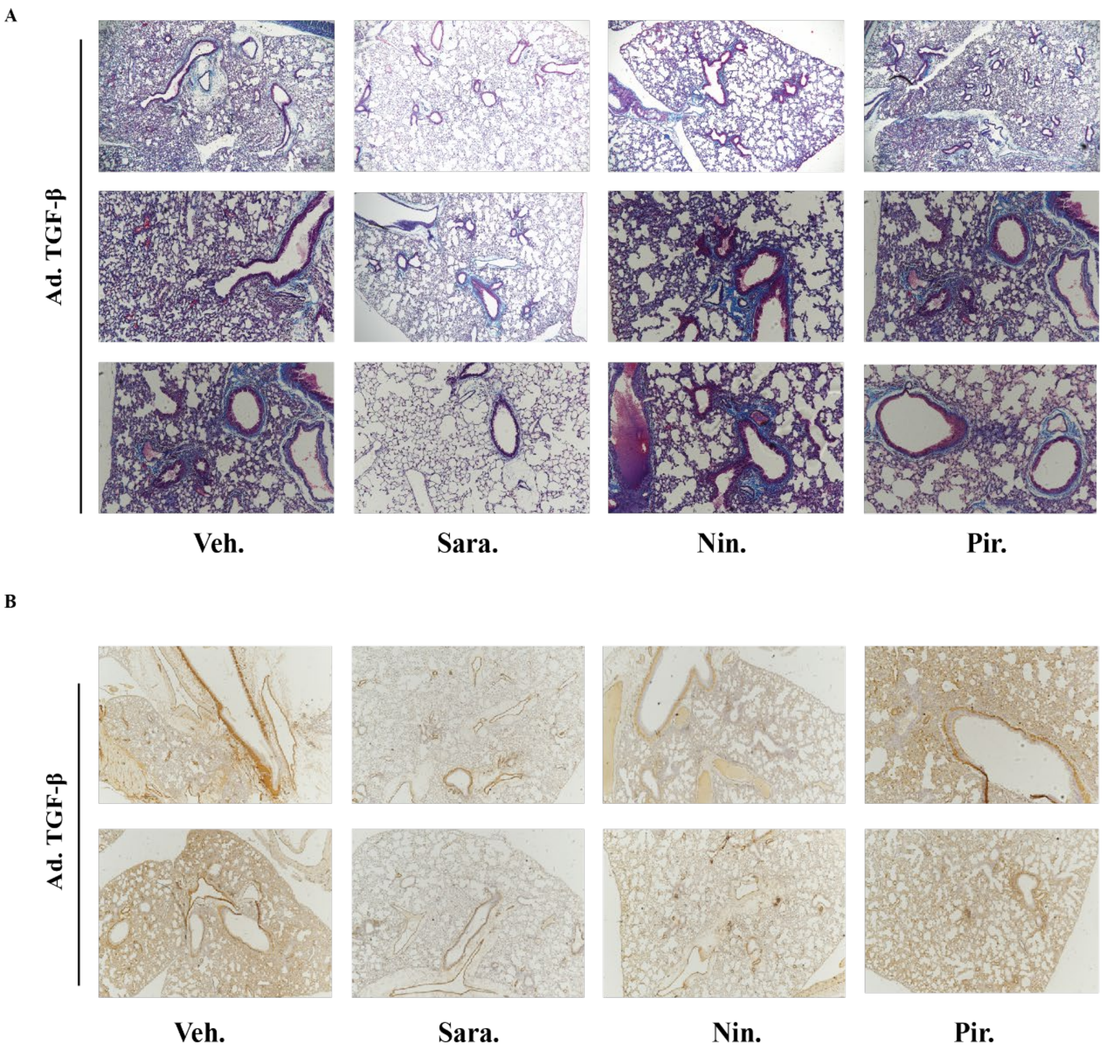


**Figure S8. Representative images of lung tissues in Ad-TGF-β mouse model.**

(A) Representative images of Masson's Trichrome staining of lung sections in the indicated groups of mice, (B) Representative images of α-SMA staining of lung sections in the indicated groups of mice.

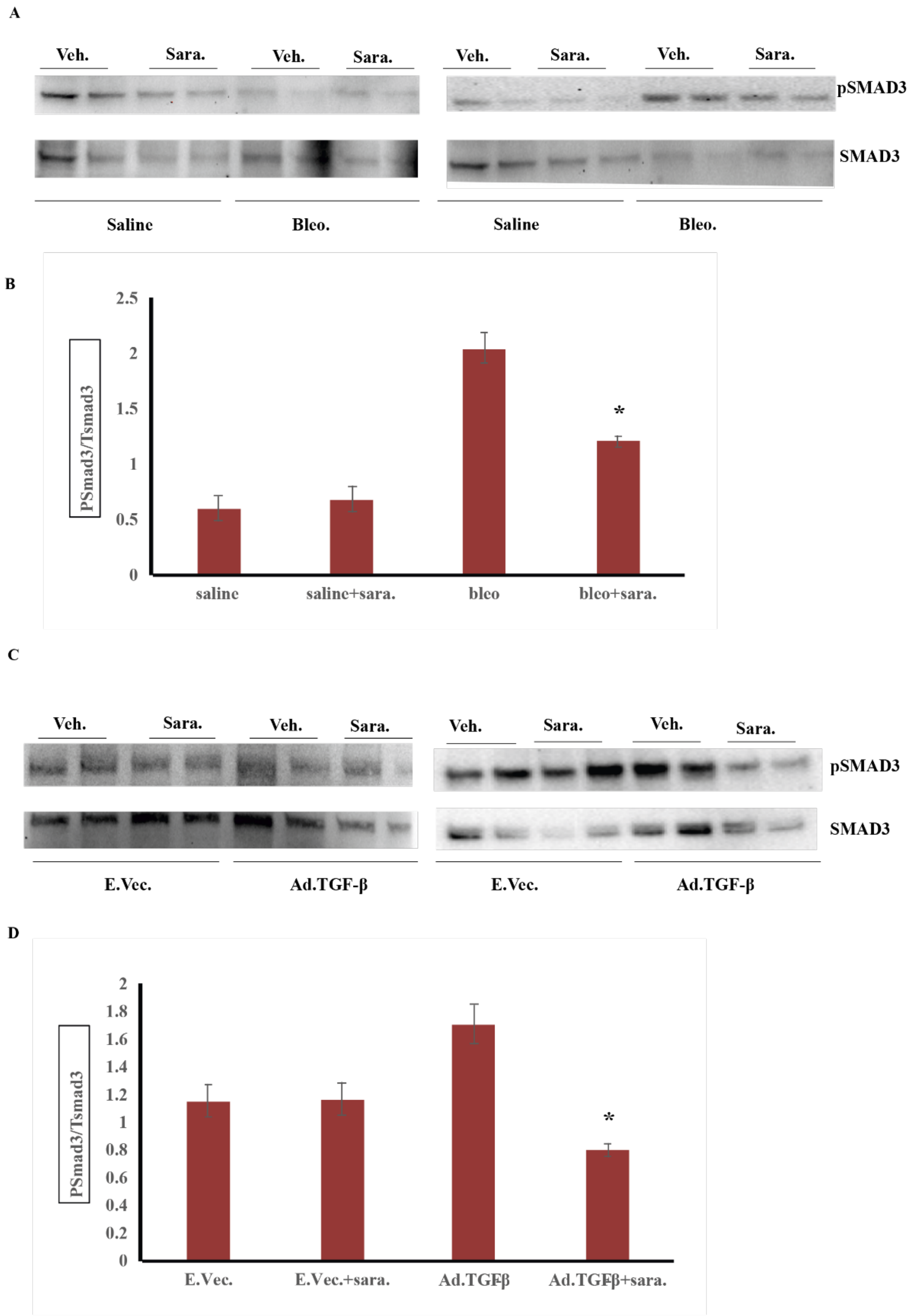
A
Veh.
Sara.
Veh.
Sara.
Veh.
Sara.
Veh.
Sara.
pSMAD3
SMAD3
Saline
Bleo.
Saline
Bleo.
B
PSmad3/Tsmad3
2.5
2
1.5
1
0.5
0
*
saline
saline+sara.
bleo
bleo+sara.
C
Veh.
Sara.
Veh.
Sara.
Veh.
Sara.
Veh.
Sara.
pSMAD3
SMAD3
E.Vec.
Ad.TGF-β
E.Vec.
Ad.TGF-β
D
PSmad3/Tsmad3
2
1.8
1.6
1.4
1.2
1
0.8
0.6
0.4
0.2
0
*
E.Vec.
E.Vec.+sara.
Ad.TGFβ
Ad.TGFβ+sara.

**Figure S9. Western blot analysis of lung tissues identified saracatinib significantly inhibits Phospho-Smad3 signaling in both animal models of pulmonary fibrosis.**

(A) Representative western blots showing saracatinib inhibits phosphorylation of Smad3 in lung tissues from Bleomycin mouse model in indicated groups. (B) Quantification of the western blots from Bleomycin mice model, data presented as means+ SEM, $*P < 0.05$.(C) Representative western blots showing saracatinib inhibits phosphorylation of Smad3 in lung tissues from Ad-TGF-β mouse model in indicated groups. (D) Quantification of the western blots from Ad-TGF-β mouse model, data presented as means+ SEM, $*P < 0.05$.

A

B

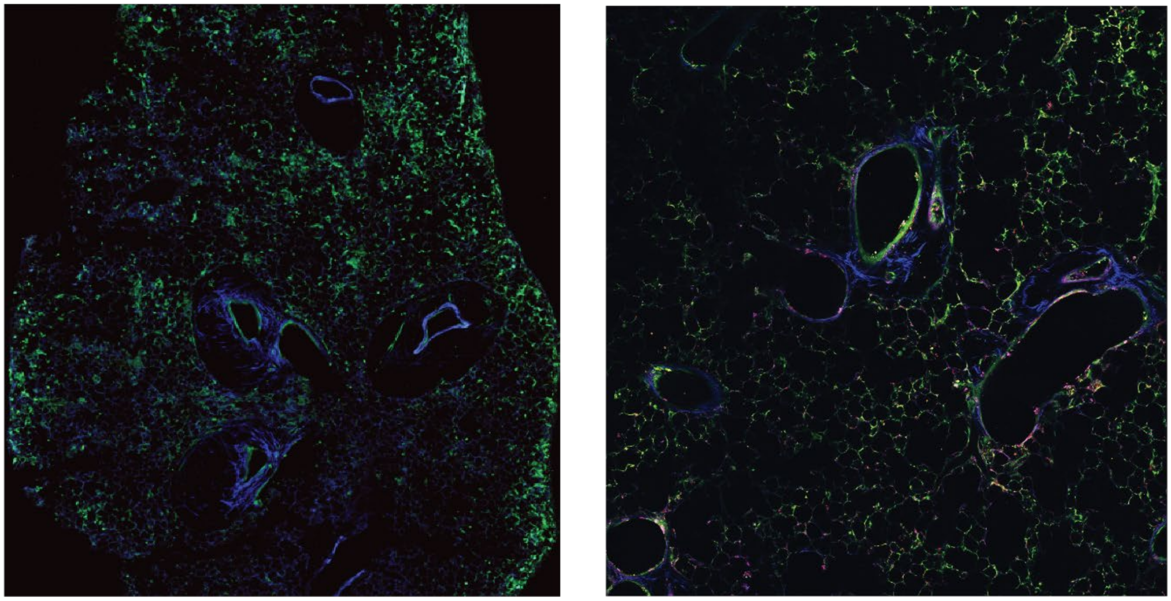

4 X

10 X

**Figure S10. LIVE/DEAD® Viability/Cytotoxicity identified the viability of PCLS after 5 days in culture.**

PCLS after 120 hours of culturing, were incubated with 2 µM Calcein AM and 2 µM Ethidium homodimer-1 (EthD-1) in 250 µm HBSS for 30 min at 37 °C. PCLS were then washed twice with HBSS and fixed with 10% buffered formalin for 30 min at RT and washed. Calcein AM (green fluorescence), live cells. Ethidium homodimer-1 (bright red fluorescence); dead cells. (A) 4X magnification (B) 10X magnification.

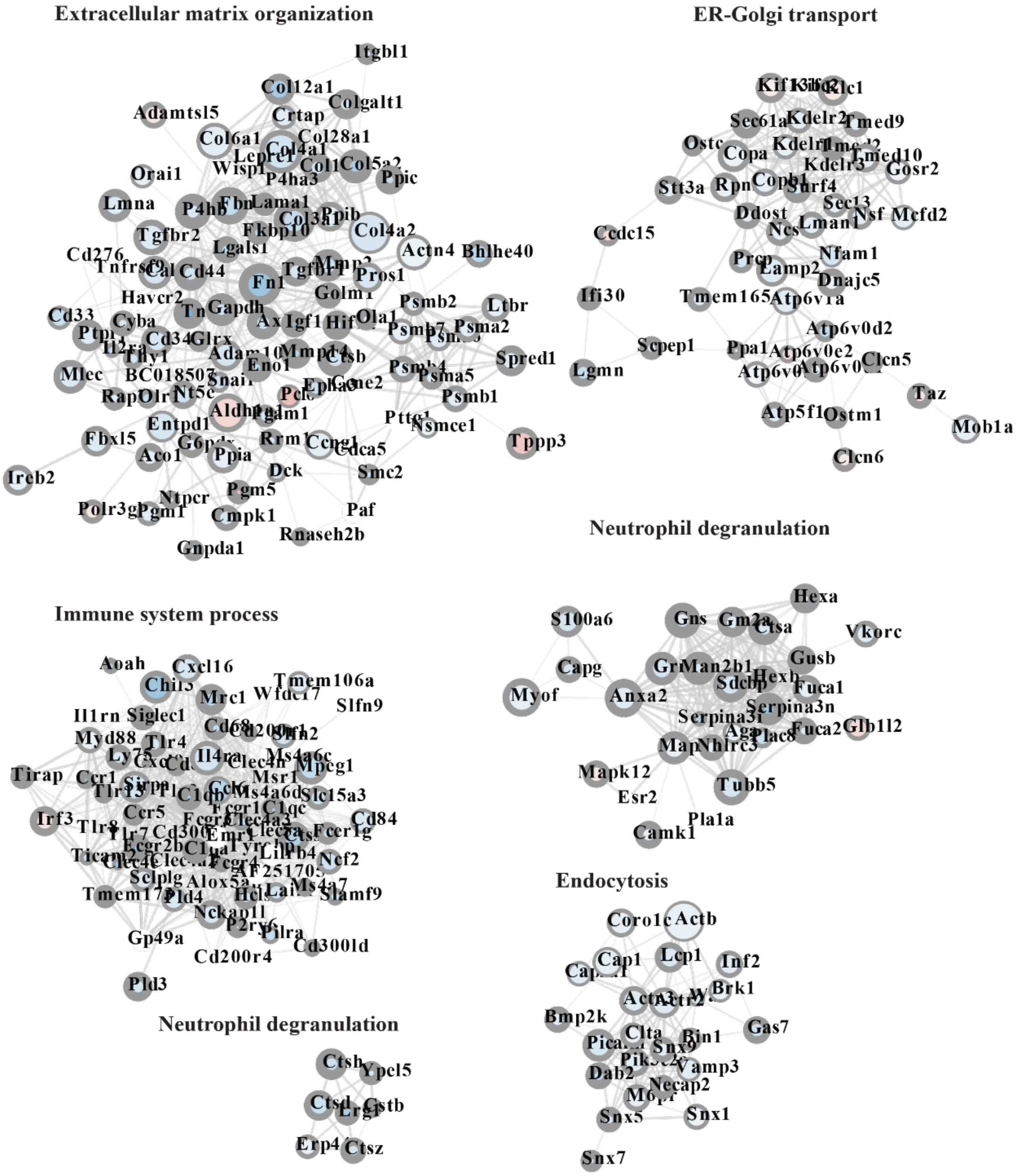

Extracellular matrix organization
ER-Golgi transport
Neutrophil degranulation
Immune system process
Neutrophil degranulation
Endocytosis
Itgbl1
Col12a1
Colgalt1
Adamtsl5
Crtap
Col6a1
Col28a1
Orai1
Ppic
Lmna
P4hb
Lama1
Tgfbr2
Fkbp10
Col4a2
Actn4
Bhlhe40
Cd276
Lgals1
Cd44
Fn1
Pros1
Havcr2
Gapdh
Golm1
Psmb2
Ltbr
Cd33
Cyba
Igf1
Psma2
Glrx
Spred1
Mlec
BC018507
Eno1
Psma5
Nt5e
Psmb1
Entpd1
Pgam1
Nsmce1
Tppp3
Fbxl5
Aco1
Ppia
Cdca5
Ireb2
Dck
Smc2
Ntpcr
Pgm5
Paf
Cmpk1
Rnaseh2b
Gnpda1
Klc1
Sec61a
Kdelr2
Tmed9
Ostc
Copa
Tmed10
Gosr2
Stt3a
Copb1
Srrf4
Sec13
Ccdc15
Ddost
Nsf
Mcfd2
Lman1
Nfam1
Dnajc5
Ifi30
Tmem165
Atp6v1a
Atp6v0d2
Scpep1
Ppa1
Atp6v0e2
Clcn5
Lgmn
Taz
Atp5f1
Ostm1
Mob1a
Clcn6
Hexa
S100a6
Gns
Vkorc
Gusb
Capg
Myof
Anxa2
Fuca1
Serpina3n
Fuca2
Glb1l2
Mapk12
Tubb5
Esr2
Pla1a
Camk1
Aoah
Cxcl16
Tmem106a
Mrc1
Slfn9
Il1rn
Siglec1
Myd88
Tlr4
Cd68
Mpeg1
Tirap
Ccr1
Msr1
Slc15a3
Ccr5
Cd84
Irf3
Tlr8
Fcer1g
Ncf2
Sclplg
Slamf9
Nckap1l
Gp49a
Pilra
Cd300ld
Cd200r4
Pld3
Ctsh
Ypel5
Ctsd
Cstb
Erp4
Ctsz
Actb
Lcp1
Inf2
Cap1
Brk1
Bmp2k
Gas7
Clta
Bin1
Dab2
Vamp3
Necap2
Snx5
Snx1
Snx7

**Figure S11. Enlarged version of Cytoscape network analysis of common differentially expressed genes shared by both the bleomycin and Ad-TGF-β mouse models that are reversed by saracatinib.**

Node size - expression level; Node color – LogFC (red is up, blue is down); node border width – negative log adj. p-value. Modules for extracellular matrix organization, immune system process, ER-golgi transport, neutrophil degranulation and endocytosis are shown.

| **Disease Signature** | **Drug Condition** | **Adjusted Score** | **FDR** |
|---|---|---|---|
| IPF- human lung – Upper lobe – Advanced – Female - GSE24206 | Low Dose MCF7 | -2.91 | $1.1 \times 10^{-19}$ |
| IPF- human lung – Upper lobe – Advanced – Male - GSE24206 | Low Dose MCF7 | -2.69 | $1.5 \times 10^{-16}$ |
| IPF- human lung fibroblasts – Slow progressing fibrosis – GSE44723 | High Dose A549 | -2.55 | $2.5 \times 10^{-11}$ |
| IPF- human lung fibroblasts – Slow progressing fibrosis – GSE44723 | Low Dose A549 | -2.38 | $1.4 \times 10^{-9}$ |
| IPF- human lung – Lower lobe – Advanced – Female - GSE24206 | Low Dose MCF7 | -2.49 | $2.6 \times 10^{-8}$ |
| IPF- human lung fibroblasts – Slow progressing fibrosis – GSE44723 | High Dose MCF7 | -1.99 | $6.1 \times 10^{-6}$ |

**Table S1. Connection between IPF and saracatinib signature.**

A significant (FDR <0.01) and negative connectivity score was detected between the SRC kinase inhibitor (saracatinib) and IPF disease signatures.

| Gene Set | # Genes in Gene Set (K) | # Genes in Overlap (k) | k/K | Description | FDR | Genes |
|---|---|---|---|---|---|---|
| **Hallmark Interferon Alpha Response** | 97 | 13 | 0.134 | Genes up regulated in response to alpha interferon proteins | 4.13 x $10^{-8}$ | PLSCR1, CASP1, IRF2, MX1, TDRD7, SP110, IFIT3, HERC6, DDX60, EPSTI1, GMPR, UBA7, GBP2 |
| **Hallmark Interferon Gamma Response** | 200 | 16 | 0.08 | Genes up regulated in response to IFNG | 3.96 x $10^{-7}$ | PLSCR1, CASP1, IRF2, MX1, TDRD7, SP110, IFIT3, HERC6, DDX60, EPSTI1, FGL2, TNFAIP6, VAMP5, IFIT1, OAS2, SECTM1 |
| **Hallmark IL2 STAT5 Signaling** | 200 | 13 | 0.0653 | Genes up regulated by STAT5 in response to IL2 stimulation | 4.26 x $10^{-5}$ | PLSCR1, FGL2, BHLHE40, NFIL3, PLIN2, PHLDA1, P2RX4, TNFRSF1B, AHR, SOCS2, IKZF2, IRF6, ADAM19 |
| **Hallmark Epithelial Mesenchymal Transition** | 200 | 13 | 0.065 | Genes defining epithelial-mesenchymal transition, as in wound healing, fibrosis and metastasis | 4.26 x $10^{-5}$ | IGFBP3, PMEPA1, TGFBR3, COL4A2, COL4A1, TNFRSF11B, TIMP3, TPM1, FSTL3, COL7A1, SERPINE2, SNTB1, CRLF1 |
| **Hallmark Hypoxia** | 200 | 13 | 0.065 | Genes up regulated in response to low oxygen levels (hypoxia) | 4.26 x $10^{-5}$ | BHLHE40, NFIL3, PLIN2, IGFBP3, CITED2, ADM, SAP30, DTNA, PDK3, MAP3K1, GCNT2, NEDD4L, CA12 |
| **Hallmark TNFα signaling via NFKB** | 200 | 13 | 0.065 | Genes regulated by NF-kB in response to TNF | 4.26 x $10^{-5}$ | TNFAIP6, BHLHE40, NFIL3, PHLDA1, PMEPA1, SMAD3, PLPP3, STAT5A, NR4A2, SNN, TRIB1, FUT4, NR4A3 |
| **Hallmark UV Response** | 144 | 11 | 0.0764 | Genes downregulated in response to ultraviolet (UV) radiation | 5.11 x $10^{-5}$ | BHLHE40, TGFBR3, CITED2, SMAD3, PLPP3, LPAR1, CACNA1A, MAGI2, GRK5, CELF2, ADGRL2 |
| **Hallmark Inflammatory Response** | 200 | 12 | 0.06 | Genes defining inflammatory response | 1.88 x $10^{-4}$ | TNFAIP6, P2RX4, TNFRSF1B, AHR, ADM, LPAR1, FZD5, P2RX7, ITGB8, TLR3, KCNJ2, NDP |
| **KEGG Pathways in Cancer** | 200 | 14 | 0.0431 | Pathways in cancer | 1.23 x $10^{-3}$ | COL4A2, COL4A1, SMAD3, STAT5A, FZD5, RAC2, WNT16, FZD4, WNT7B, MITF, HGF, FGF7, RASSF5, EPAS1 |
| **KEGG Axon Guidance** | 129 | 8 | 0.062 | Axon guidance | 4.18 x $10^{-3}$ | RAC2, FES, EFNA5, SEMA6C, SEMA4B, SEMA3D, NGEF, SEMA3F |

**Table S2. Enrichment analysis of significantly differentially expressed genes.**

Saracatinib induced alterations in numerous pathways identified from the Hallmark and KEGG databases, with IFNα, IFNγ, EMT, and inflammatory responses among the top pathways (FDR <

0.05) in NHLFs incubated with saracatinib followed by TGF-β treatment. The top 10 pathways are shown (significant at FDR < 0.05).
